# *Plasmodium falciparum* K13 mutations in Africa and Asia present varying degrees of artemisinin resistance and an elevated fitness cost in African parasites

**DOI:** 10.1101/2021.01.05.425445

**Authors:** Barbara H. Stokes, Kelly Rubiano, Satish K. Dhingra, Sachel Mok, Judith Straimer, Nina F. Gnädig, Jade R. Bath, Ioanna Deni, Kurt E. Ward, Josefine Striepen, Tomas Yeo, Leila S. Ross, Eric Legrand, Frédéric Ariey, Clark H. Cunningham, Issa M. Souleymane, Adama Gansané, Romaric Nzoumbou-Boko, Claudette Ndayikunda, Abdunoor M. Kabanywanyi, Aline Uwimana, Samuel J. Smith, Olimatou Kolley, Mathieu Ndounga, Marian Warsame, Rithea Leang, François Nosten, Timothy J.C. Anderson, Philip J. Rosenthal, Didier Ménard, David A. Fidock

## Abstract

The emergence of artemisinin (ART) resistance in *Plasmodium falciparum* parasites has led to increasing rates of treatment failure with first-line ART-based combination therapies (ACTs) in Southeast Asia. In this region, select mutations in K13 can result in delayed parasite clearance rates *in vivo* and enhanced survival in the ring-stage survival assay (RSA) *in vitro*. Our genotyping of 3,299 *P. falciparum* isolates across 11 sub-Saharan countries reveals the continuing dominance of wild-type K13 and confirms the emergence of a K13 R561H variant in Rwanda. Using gene editing, we provide definitive evidence that this mutation, along with M579I and C580Y, can confer variable degrees of *in vitro* ART resistance in African *P. falciparum* strains. C580Y and M579I were both associated with substantial fitness costs in African parasites, which may counter-select against their dissemination in high-transmission settings. We also report the impact of multiple K13 mutations, including the predominant variant C580Y, on RSA survival rates and fitness in multiple Southeast Asian strains. No change in ART susceptibility was observed upon editing point mutations in ferrodoxin or mdr2, earlier associated with ART resistance in Southeast Asia. These data point to the lack of an evident biological barrier to mutant K13 mediating ART resistance in Africa, while identifying their detrimental impact on parasite growth.

## Introduction

Despite recent advances in chemotherapeutics, diagnostics and vector control measures, malaria continues to exert a significant impact on human health. In 2019, cases were estimated at 229 million, resulting in 409,000 fatal outcomes, primarily in Sub-Saharan Africa as a result of *Plasmodium falciparum* infection (WHO, 2020). This situation is predicted to rapidly worsen as a result of the ongoing SARS-CoV-2 pandemic that has crippled malaria treatment and prevention measures (Sherrard-Smith *et al.*, 2020). Absent an effective vaccine, malaria control and elimination strategies are critically reliant on the continued clinical efficacy of first-line artemisinin-based combination therapies (ACTs). These ACTs pair fast-acting artemisinin (ART) derivatives with partner drugs such as lumefantrine, amodiaquine, mefloquine or piperaquine (PPQ). ART derivatives can reduce the biomass of drug-sensitive parasites by up to 10,000-fold within 48 h (the duration of one intra-erythrocytic developmental cycle); however, these derivatives are rapidly metabolized *in vivo*. Longer-lasting albeit slower-acting partner drugs are co-administered to reduce the selective pressure for ART resistance and to clear residual parasitemias.

Nonetheless, *P. falciparum* partial resistance to ART derivatives has spread throughout Southeast Asia (SEA), having first emerged a decade ago in western Cambodia (Dondorp *et al.*, 2009; Noedl *et al.*, 2009; Ariey *et al.*, 2014; Imwong *et al.*, 2020). Clinically, partial ART resistance manifests as delayed clearance of circulating asexual blood stage parasites following treatment with an ACT. The accepted threshold for resistance is a parasite clearance half-life (the time required for the peripheral blood parasite density to decrease by 50%) of >5.5 h. Sensitive parasites are typically cleared in <2-3 h (WHO, 2019). Partial resistance can also be evidenced as parasite-positive blood smears on day three post initiation of treatment. *In vitro,* ART resistance manifests as increased survival of cultured ring-stage parasites exposed to a 6 h pulse of 700 nM dihydroartemisinin (DHA, the active metabolite of all ARTs used clinically) in the ring-stage survival assay (RSA) (Witkowski *et al.*, 2013; Ariey *et al.*, 2014). Recently, ART-resistant strains have also acquired resistance to PPQ, which is widely used in SEA as a partner drug in combination with DHA. Failure rates following DHA-PPQ treatment now exceed 50% in parts of Cambodia, Thailand and Vietnam (van der Pluijm *et al.*, 2019).

*In vitro* selections, supported by clinical epidemiological data, have demonstrated that ART resistance is primarily determined by mutations in the beta-propeller domain of the *P. falciparum* Kelch protein K13 (Ariey *et al.*, 2014; Ashley *et al.*, 2014; MalariaGEN, 2016; Menard *et al.*, 2016; Siddiqui *et al.*, 2020). Recent evidence suggests that these mutations result in reduced endocytosis of host-derived hemoglobin and thereby decreased release of the ART-activating moiety Fe^2+^-heme, thus reducing ART potency (Yang *et al.*, 2019; Birnbaum *et al.*, 2020). Mutations in other genes including ferredoxin *(fd)* and multidrug resistance protein 2 *(mdr2)* have also been associated with ART resistance in K13 mutant parasites, suggesting that they either contribute to a multigenic basis of resistance or fitness or serve as genetic markers of founder populations (Miotto *et al.*, 2015).

In SEA, the most prevalent K13 mutation is C580Y, which associates with delayed clearance *in vivo* (Ariey *et al.*, 2014; Ashley *et al.*, 2014; MalariaGEN, 2016; Menard *et al.*, 2016; Imwong *et al.*, 2017). This mutation also mediates ART resistance *in vitro,* as demonstrated by RSAs on gene-edited parasites (Ghorbal *et al.*, 2014; Straimer *et al.*, 2015; Uwimana *et al.*, 2020). Other studies have since demonstrated the emergence of almost 200 K13 mutations both in SEA and other malaria-endemic regions, including India, the Guiana Shield and the western Pacific (MalariaGEN, 2016; Menard *et al.*, 2016; Das *et al.*, 2019; Group, 2019; Mathieu *et al.*, 2020; Miotto *et al.*, 2020). Aside from C580Y, only a handful of K13 mutations (N458Y, M476I, Y493H, R539T, I543T and R561H) have been validated by gene-editing experiments as conferring resistance *in vitro* (Straimer *et al.*, 2015; Siddiqui *et al.*, 2020). Multiple other mutations have been associated with the clinical delayed clearance phenotype and have been proposed as candidate markers of ART resistance (Group, 2019; WHO, 2019). Here, we define the role of a panel of K13 mutations found in field isolates, and address the key question of whether these mutations can confer resistance in African strains. We include the K13 R561H mutation, earlier associated with delayed parasite clearance in SEA (Ashley *et al.*, 2014; Phyo *et al.*, 2016), and very recently identified at 7-12% prevalence in certain districts in Rwanda (Uwimana *et al.*, 2020; Bergmann *et al.*, 2021). This study also enabled us to assess the impact of the parasite genetic background, including ferrodoxin and mdr2 as proposed secondary determinants of resistance, on *in vitro* phenotypes. Our results show that K13 mutations can impart ART resistance across multiple Asian and African strains, at levels that vary widely depending on the mutation and the parasite genetic background. Compared to K13 mutant Asian parasites, we observed stronger *in vitro* fitness costs in K13-edited African strains, which might predict a slower dissemination of ART resistance in high-transmission African settings.

## Results

### Non-synonymous K13 mutations are present at low frequencies in Africa

To examine the status of K13 mutations across Africa, we sequenced the beta-propeller domain of this gene in 3,299 isolates from 11 malaria-endemic African countries, including The Gambia, Sierra Leone, and Burkina Faso in West Africa; Chad, Central African Republic, Republic of the Congo, and Equatorial Guinea in Central Africa; and Burundi, Tanzania, Rwanda, and Somalia in East Africa. Samples were collected between 2011 and 2019, with most countries sampled across multiple years.

Of all samples, 99% (3,220) were K13 WT, i.e. they matched the 3D7 (African) reference sequence or harbored a synonymous (non-coding) mutation. For individual countries, the percentage of K13 WT samples ranged from 95% to 100% (**Figure 1; Table S1**). In total, we identified 36 unique non-synonymous mutations in K13. Only two of these non-synonymous mutations have been validated as resistance mediators in the Southeast Asian Dd2 strain: the M476I mutation initially identified from long-term ART selection studies and the R561H mutation observed in Rwanda (Ariey *et al.*, 2014; Straimer *et al.*, 2015; Uwimana *et al.*, 2020).

**Figure 1.**
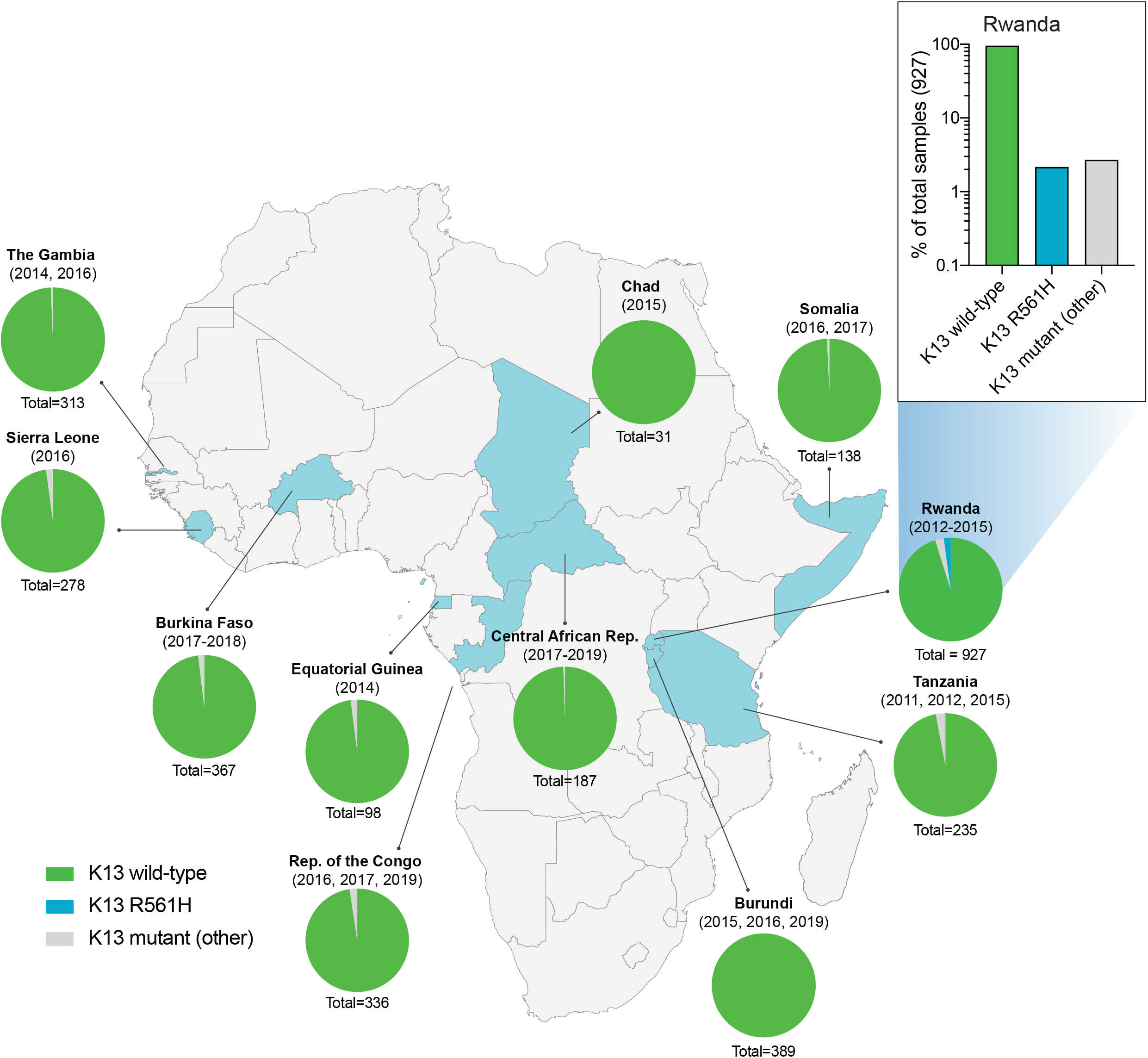
Frequency and distribution of *K13* alleles in 11 African countries. Pie charts representing the proportions of sequenced samples per country that harbor the K13 wild-type (WT; 3D7 reference) sequence, the R561H variant (the most commonly identified mutation, unique to Rwanda; see inset), or other less frequent non-synonymous K13 mutations. Sample sizes and years of sample collection are indicated. Mutations and numbers of African samples sequenced per country are listed in **Table S1**.

Of the 36 non-synonymous mutations, only two were present in >6 samples: R561H (n=20, unique to Rwanda, sampled from 2012 to 2015), and A578S (n=10; observed in four countries across multiple years). Previously A578S was shown not to confer *in vitro* resistance in Dd2 (Menard *et al.*, 2016). R561H accounted for 44% of mutant samples and 7% of all samples in the set of 927 Rwandan genotyped isolates (Uwimana *et al.*, 2020).

### The K13 R561H, M579I and C580Y mutations can confer *in vitro* artemisinin resistance in African parasites

To test whether R561H can mediate ART resistance in African strains, we developed a CRISPR/Cas9-mediated *K13* editing strategy (**Figure S1**) to introduce this mutation into 3D7 and F32 parasites. On the basis of whole-genome sequence analysis of African isolates, 3D7 was recently shown to segregate phylogenetically with parasites from Rwanda (Ariey *et al.*, 2014; Uwimana *et al.*, 2020). F32 was derived from an isolate from Tanzania (Witkowski *et al.*, 2010). We also tested the C580Y mutation that predominates in SEA, as well as the M579I mutation identified in a *P. falciparum*-infected individual in Equatorial Guinea who displayed delayed parasite clearance following ACT treatment (Lu *et al.*, 2017). The positions of these residues are highlighted in the K13 beta-propeller domain structure shown in **Figure S2**. For 3D7, F32 and other lines used herein, the geographic origins and genotypes at drug resistance loci are described in **Table 1** and **Table S2**. All parental lines were cloned by limiting dilution prior to transfection. Edited parasites were identified by PCR and Sanger sequencing, and cloned.

**Table 1.**
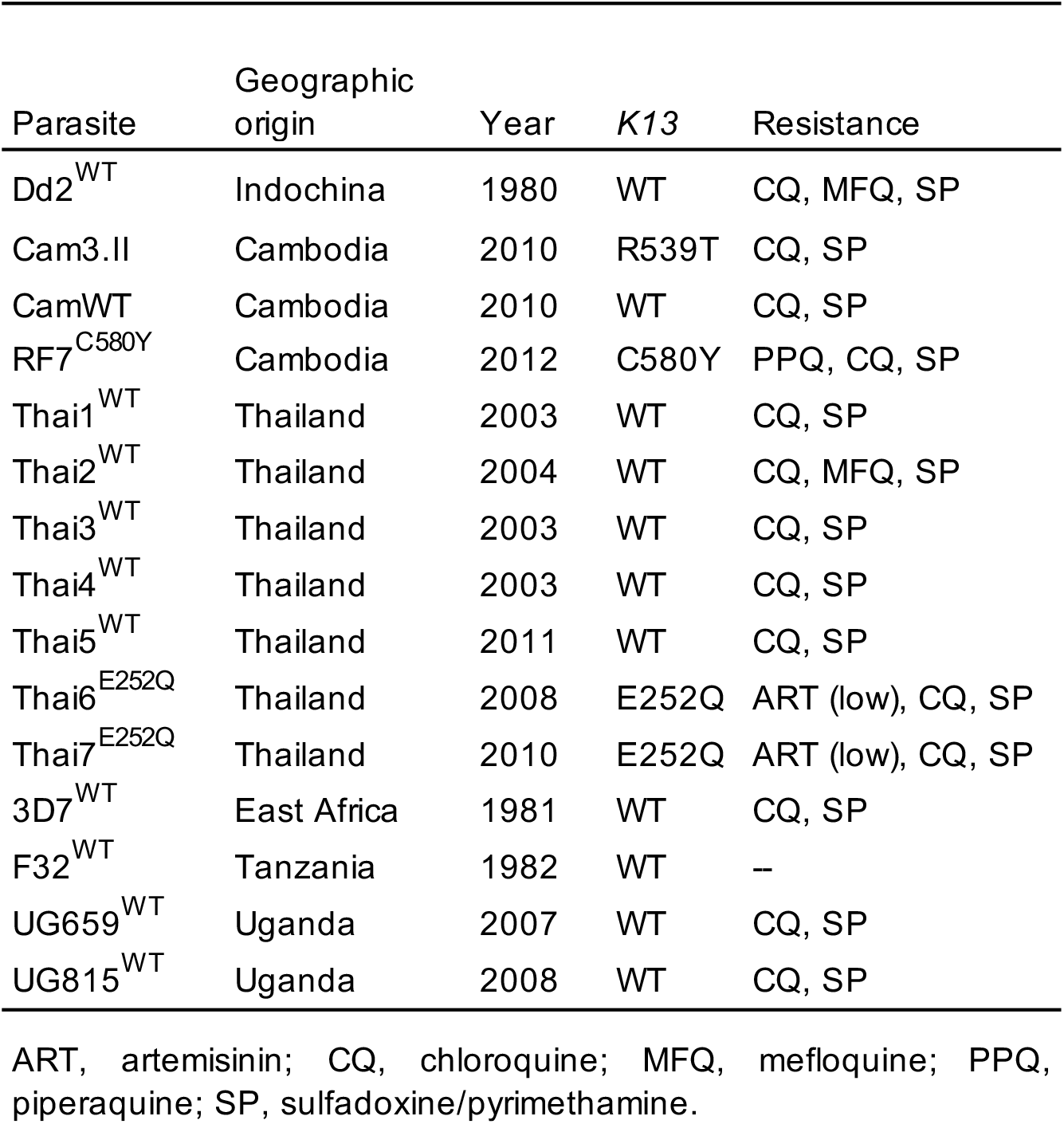
*Plasmodium falciparum* lines employed herein.

RSAs, used to measure *in vitro* ART susceptibility, revealed a wide range of mean survival values for K13 mutant lines. For 3D7 parasites, the highest RSA survival rates were observed with 3D7^R561H^ parasites, which averaged 6.6% RSA survival. For the 3D7^M579I^ and 3D7^C580Y^ lines, mean RSA survival rates were both 4.8%, a 3 to 4-fold increase relative to the 3D7^WT^ line. No elevated RSA survival was seen in a 3D7 control line (3D7^ctrl^) that expressed only the silent shield mutations used at the guide RNA cut site (**Figure 2A; Table S3**).

**Figure 2.**
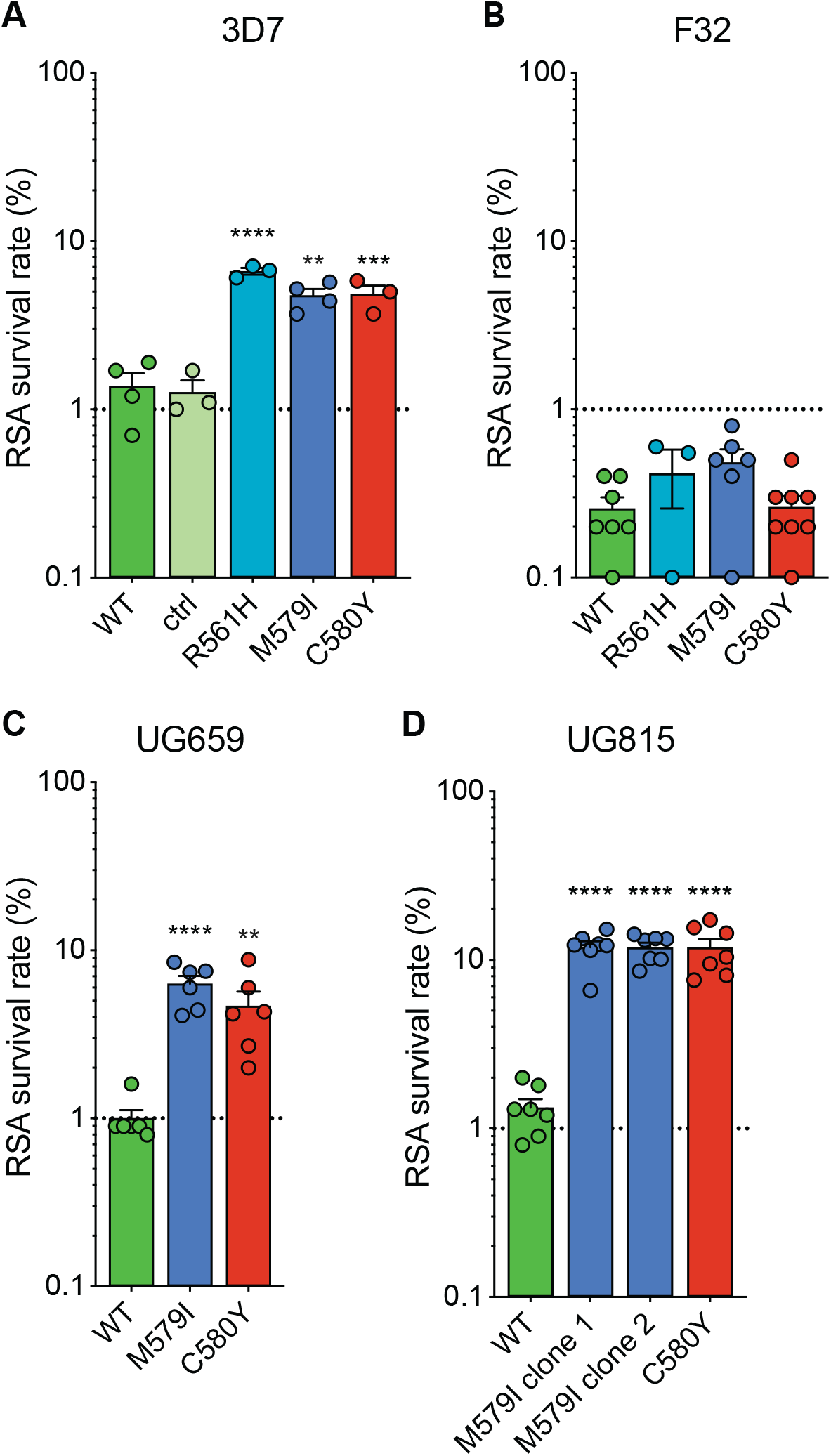
Gene-edited African parasites acquire variable degrees of *in vitro* ART resistance conferred by mutant K13. (A-D) RSA survival rates for (A) 3D7 (Africa), (B) F32 (Tanzania), (C) UG659 (Uganda), or (D) UG815 (Uganda) K13 wild-type parental lines and CRISPR/Cas9-edited K13 R561H, M579I or C580Y mutant clones. Parental lines are described in **Table 1** and **Table S2**. For 3D7, we also included a K13 wild-type control (ctrl) line harboring silent-binding site mutations at the K13 gRNA cut site. Results show the percentage of early ring-stage parasites (0-3 hpi) that survived a 6 h pulse of 700 nM DHA, relative to DMSO-treated parasites assayed in parallel. Percent survival values are shown as means ± SEM (detailed in **Table S3**). Results were obtained from 3 to 8 independent experiments, each performed in duplicate. *P* values were determined by unpaired *t* tests and were calculated for mutant lines relative to the isogenic line expressing WT *K13*. ** *P*<0.01; *** *P*<0.001; **** *P*<0.0001.

In contrast to results with 3D7, the introduction of K13 mutations into F32^WT^ parasites yielded almost no increase in RSA survival. Mean RSA survival rates were 0.5%, 0.5% and 0.3% for F32^R561H^, F32^M579I^ and F32^C580Y^ parasites, respectively, compared to 0.3% for F32^WT^ (**Figure 2B**). Previously we reported that introduction of M476I into F32 parasites resulted in a modest gain of resistance (mean survival of 1.7%) while this same mutation conferred RSA survival levels of ~10% in edited Dd2 parasites (Straimer *et al.*, 2015). These data suggest that while K13 mutations differ substantially in the level of resistance that they impart, there is an equally notable contribution of the parasite genetic background.

We next introduced M579I and C580Y into cloned Ugandan isolates UG659 and UG815. Editing of both mutations into UG659 yielded moderate RSA survival rates (means of 6.3% and 4.7% for UG659^M579I^ or UG659^C580Y^ respectively, vs. 1.0% for UG659^WT^; **Figure 2C**). These values resembled our results with 3D7. Strikingly, introducing K13 M579I or C580Y into UG815 yielded the highest rates of *in vitro* resistance, with mean survival levels reaching ~12% in both UG815^M579I^ and UG815^C580Y^. These results were confirmed in a second independent clone of UG815^M579I^ (**Figure 2D**). M579I and C580Y also conferred equivalent levels of resistance in edited Dd2 parasites (RSA survival rates of 4.0% and 4.7%, respectively; **Table S3**). These data show that mutant K13-mediated ART resistance in African parasites can be achieved, in some but not all strains, at levels comparable to or above those seen in Southeast Asian parasites.

### The K13 C580Y and M579I mutations are associated with an *in vitro* fitness defect in African parasites

To examine the relation between resistance and fitness in African parasites harboring K13 mutations, we developed an *in vitro* fitness assay that uses quantitative real-time PCR (qPCR) for allelic discrimination. Assays were conducted with the eight pairs of K13 WT and either C580Y or M579I isogenic parasites used for RSAs, namely 3D7^WT^ + either 3D7^M579I^ or 3D7^C580Y^; F32^WT^ + either F32^M579I^ or F32^C580Y^; UG659^WT^ + either UG659^M579I^ or UG659^C580Y^; and UG815^WT^ + either UG815^M579I^ or UG915^C580Y^.

Assays were initiated with tightly synchronized trophozoites, mixed in 1:1 ratios of WT to mutant isogenic parasites, and cultures were maintained over a period of 40 days (~20 generations of asexual blood stage growth). Cultures were sampled every four days for genomic DNA (gDNA) preparation and qPCR analysis. TaqMan probes specific to the *K13* WT or mutant (M579I or C580Y) alleles were used to quantify the proportion of each allele.

Results showed that both K13 mutations (M579I or C580Y) conferred a significant fitness defect in all backgrounds tested, with the percentage of the *K13* mutant allele declining over time in all co-cultures (**Figure 3; Table S4**). This fitness defect varied between parasite backgrounds. To quantify this impact, we calculated the fitness cost, which represents the percent reduction in growth rate per 48 h generation of the test line compared to the WT isogenic comparator. The fitness cost was calculated using day 0 and day 32 as start and end points, respectively, as these yielded the most consistent slopes across lines and time series. For 3D7 parasites, the fitness cost was 8.9% and 6.9% for both the M579I and C580Y mutations, respectively (**Figure 3A**). In F32 and UG659 parasites, the fitness cost for the M579I mutation was slightly higher (4.3% and 5.8%) than for C580Y (2.8% and 1.9%; **Figure 3B, C**). The largest growth defects for both mutations were seen in the UG815 line (**Figure 3D**), with fitness cost values for the M579I and C580Y mutations both being 12.0%. A comparison of data across these four African strains revealed that high RSA survival rates were generally accompanied by high fitness costs, with M579I mostly having a more detrimental fitness impact than C580Y (**Figure 3E, 3F**).

**Figure 3.**
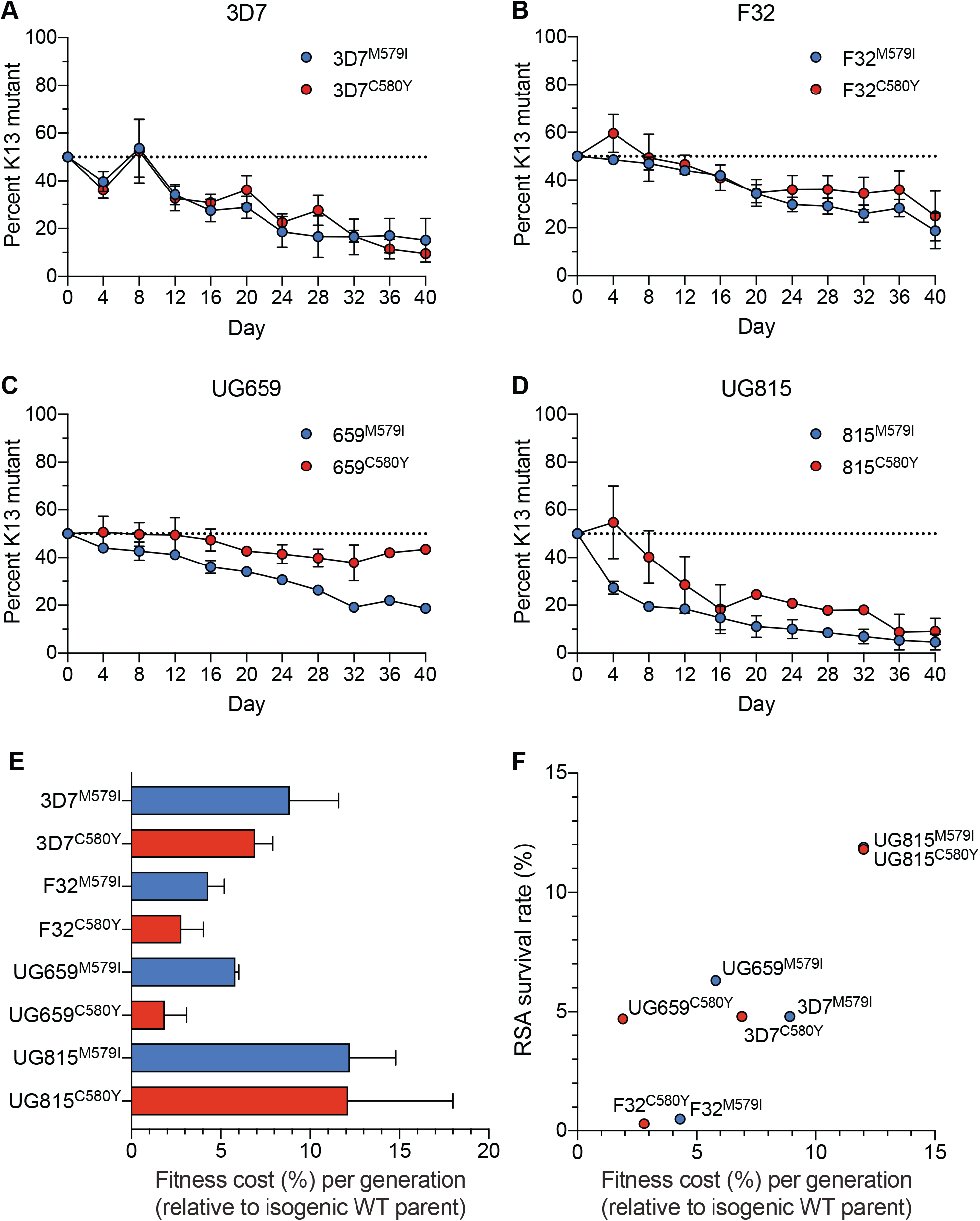
K13 M579I and C580Y mutations cause *in vitro* growth defects in gene-edited African parasites. (A-D) Percentage of mutant allele relative to the WT allele over time in (A) 3D7, (B) F32, (C) UG659, and (D) UG815 parasite cultures in which K13 M579I or C580Y mutant clones were co-cultured as 1:1 mixtures with isogenic K13 WT controls over a period of 40 days. Percentages of mutant and WT alleles in culture over time were determined by TaqMan qPCR-based allelic discrimination, and normalized to 50% on Day 0. Results, shown as means ± SEM, were obtained from 2 to 5 independent experiments, each performed in duplicate. Values are provided in **Table S4**. (E) The percent reduction in growth rate per 48 h generation, termed the fitness cost, is presented as mean ± SEM for each mutant line relative to its isogenic wild-type comparator. (F) Fitness costs for mutant lines and isogenic wild-type comparators are plotted relative to RSA survival values for the same lines.

### The K13 C580Y mutation has swept rapidly across Cambodia, displacing other K13 variants

We next examined the spatiotemporal distribution of *K13* alleles in Cambodia, the epicenter of ART resistance in SEA. In total, we sequenced the K13 propeller domains of 3,327 parasite isolates collected from western, northern, eastern and southern provinces of Cambodia (**Figure S3**). Samples were collected between 2001 and 2017, except for the southern region where sample collection was initiated in 2010. In sum, 19 nonsynonymous polymorphisms in K13 were identified across all regions and years. Of these, only three were present in >10 samples, Y493H (n=83), R539T (n=87) and C580Y (n=1,915). Each of these mutations was previously shown to confer ART resistance *in vitro* (Straimer *et al.*, 2015). Rarer mutations included A418V, I543T, P553L, R561H, P574L, and D584V (**Figure 4; Table S5**).

**Figure 4.**
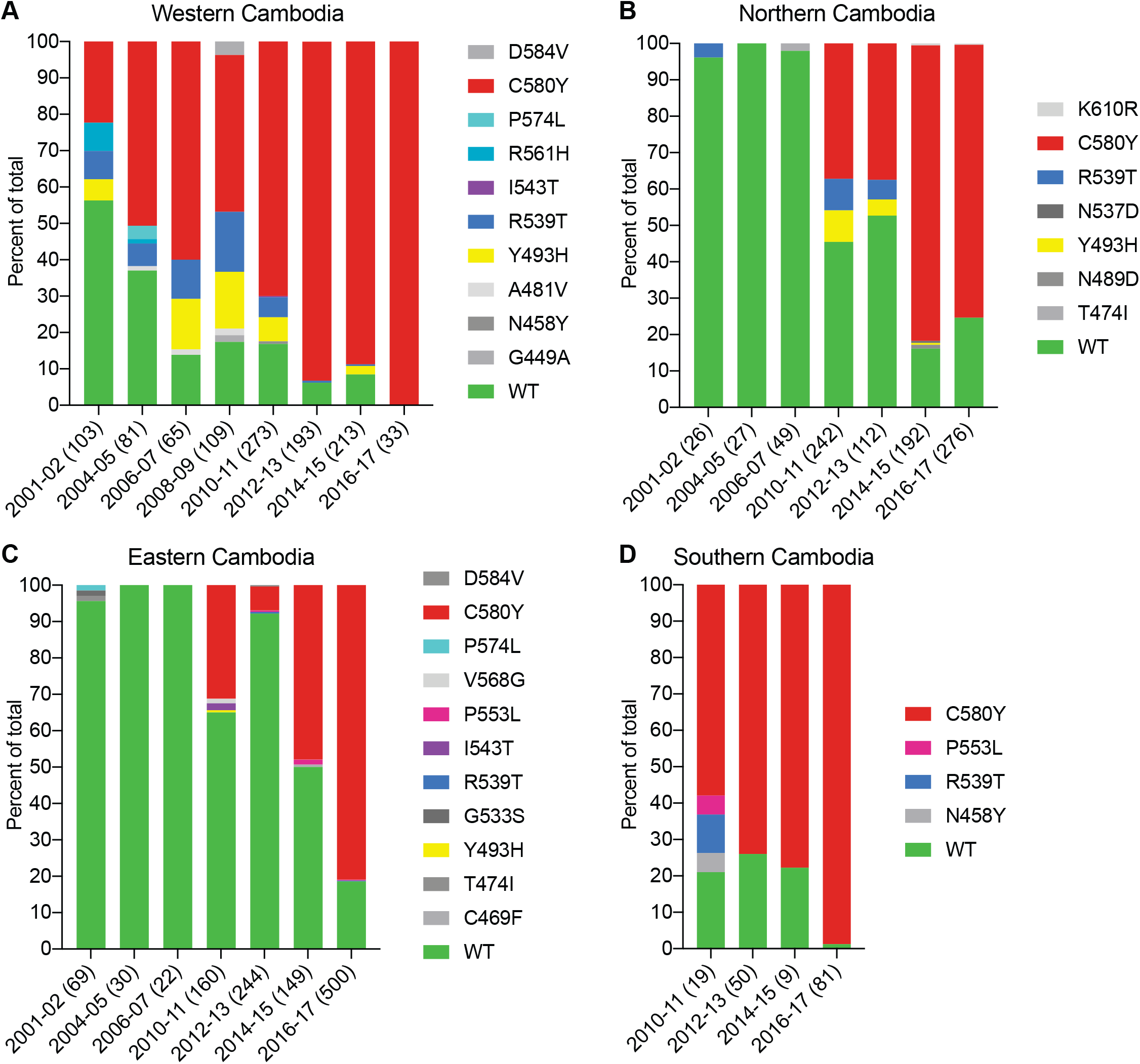
The *K13* C580Y allele has progressively outcompeted all other alleles in Cambodia. (A-D) Stacked bar charts representing the percentage of sequenced samples expressing the *K13* wild-type allele or individual variants, calculated based on the total number of samples (listed in parentheses) for a given period. Sample collection was segregated into four regions in Cambodia (detailed in **Figure S3**). All K13 mutant samples harbored a single non-synonymous nucleotide polymorphism. Mutations and numbers of Cambodian samples sequenced per region/year are listed in **Table S5**.

This analysis revealed a significant proportion of K13 WT parasites in the early 2000s, particularly in northern and eastern Cambodia, where 96% of isolates in 2001-2002 had the WT *K13* sequence (**Figure 4**). In western Cambodia, where ART resistance first emerged (Dondorp *et al.*, 2009; Noedl *et al.*, 2009), the WT allele percentage in 2001-2002 had already fallen to 56%. This is striking given that delayed parasite clearance following ACT or artesunate treatment was first documented in 2008-2009 (Noedl *et al.*, 2008; Noedl *et al.*, 2009).

In all four regions, the frequency of the WT allele declined substantially over time and the diversity of mutant alleles contracted, with nearly all WT and non-K13 C580Y mutant parasites being replaced by parasites harboring the C580Y mutation (**Figure 4**). This effect was particularly pronounced in the west and the south, where the prevalence of C580Y in 2016-17 effectively attained 100%, increasing from 22% and 58% respectively in the initial sample sets (**Figure 4A, D**). In northern and eastern Cambodia, C580Y also outcompeted all other mutant alleles; however, 19-25% of parasites remained K13 WT in 2016-17 (**Figure 4B, C**). These data show the very rapid dissemination of K13 C580Y across Cambodia.

### Southeast Asian K13 mutations associated with delayed parasite clearance differ substantially in their ability to confer artemisinin resistance *in vitro*

Given that most K13 polymorphisms present in the field have yet to be characterized *in vitro*, we selected a panel of mutations to test by gene editing, namely E252Q, F446I, P553L, R561H and P574L (**Figure S2**). The F446I mutation is the predominant mutation in the Myanmar-China border region. P553L, R561H and P574L have each been shown to have multiple independent origins throughout SEA (Menard *et al.*, 2016), and were identified at low frequencies in our sequencing study in Cambodia (**Figure 4**). Lastly, the E252Q mutation was formerly prevalent on the Thai-Myanmar border, and, despite its occurrence upstream of the beta-propeller domain, has been associated with delayed parasite clearance *in vivo* (Anderson *et al.*, 2017; Cerqueira *et al.*, 2017; Group, 2019).

Zinc-finger nuclease-or CRISPR/Cas9-based gene edited lines expressing K13 E252Q, F4461, P553L, R561H or P574L were generated in Dd2 or Cam3.II lines expressing WT K13 (Dd2^WT^ or Cam3.II^WT^) and recombinant parasites were cloned. Early ring-stage parasites were then assayed for their ART susceptibility using the RSA. For comparison, we included published Dd2 and Cam3.II lines expressing either K13 C580Y (Dd2^C580Y^ and Cam3.II^C580Y^) or R539T (Dd2^R539T^ and the original parental line Cam3.II^R539T^) (Straimer *et al.*, 2015), as well as control lines expressing only the guide-specific silent shield mutations (Dd2^ctrl^ and Cam3.II^ctrl^).

Both the P553L and R561H mutations yielded mean RSA survival rates comparable to C580Y (4.6% or 4.3% RSA survival for Dd2^P553L^ or Dd2^R561H^, respectively, vs 4.7% for Dd2^C580Y^; **Figure 5A; Table S6**). F446I and P574L showed only modest increases in survival relative to the WT parental line (2.0% and 2.1% for Dd2^F446I^ and Dd2^P574L^, respectively, vs 0.6% for Dd2^WT^). No change in RSA survival relative to Dd2^WT^ was observed for the Dd2^E252Q^ line. The resistant benchmark Dd2^R539T^ showed a mean RSA survival level of 20.0%.

**Figure 5.**
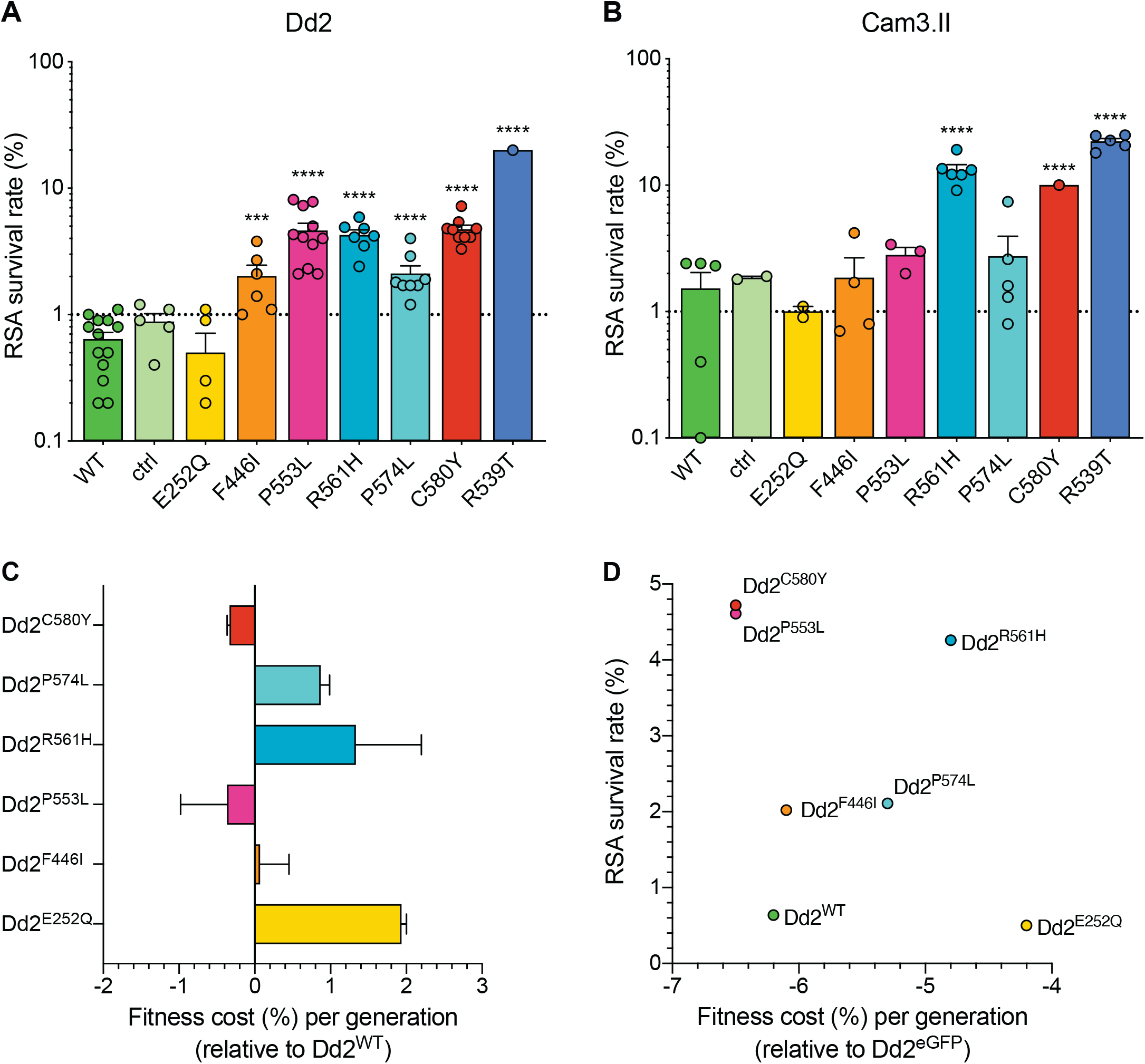
Southeast Asian K13 mutations yield elevated RSA survival and minor impacts on *in vitro* growth in gene-edited parasite lines. (A, B) RSA survival rates for Dd2 (Indochina) and Cam3.II (Cambodia) *P. falciparum* parasites expressing wild-type or mutant K13. Gene-edited parasites were generated using CRISPR/Cas9 or zinc-finger nucleases (ZFNs). Control (ctrl) lines express silent-binding site mutations at the *K13* gRNA cut site. Unedited parental lines are described in **Table 1** and **Table S2**. Results show the percentage of early ring-stage parasites (0-3 hpi) that survived a 6 h pulse of 700 nM DHA, relative to DMSO-treated parasites processed in parallel. Percent survival values are shown as means ± SEM (detailed in **Table S6**). Results were obtained from 3 to 13 independent experiments, each performed in duplicate. *P* values were determined by unpaired *t* tests and were calculated for mutant lines relative to the isogenic line expressing WT *K13*. *** *P*<0.001; **** *P*<0.0001. (C) Percent reductions in growth rate per 48 h generation, expressed as the fitness cost, for each Dd2 mutant line relative to the Dd2^WT^ line. Fitness costs were determined by co-culturing a Dd2^eGFP^ reporter line with either the Dd2 K13 wild-type parental line (Dd2^WT^) or gene-edited K13 mutant lines. Co-cultures were maintained for 20 days and percentages of eGFP^+^ parasites were determined by flow cytometry (see **Table S7 and Figure S4A**). Fitness costs were calculated relative to the Dd2^eGFP^ reporter line (**Figure S4B**) and are shown here normalized to the Dd2^WT^ line. Mean ± SEM values were obtained from three independent experiments, each performed in triplicate. (D) Fitness costs for K13 WT or mutant lines, relative to the Dd2^eGFP^ line, plotted against their corresponding RSA survival values.

In contrast to Dd2, editing of the F446I, P553L and P574L mutations into the Cambodian Cam3.II parasite background did not result in a statistically significant increase in survival relative to K13 WT Cam3.II^WT^ parasites, in part because the background survival rate of the Cam3.II^WT^ line was higher than for Dd2^WT^. All survival values were <3%, contrasting with the Cam3.II^R539T^ parental line that expresses the R539T mutation (~20% mean survival; **Figure 5B; Table S6**). The E252Q mutation did not result in elevated RSA survival in the Cam3.II background, a result also observed with Dd2. Nonetheless, ART resistance was apparent upon introducing the R561H mutation into Cam3.II parasites, whose mean survival rates exceeded the Cam3.II^C580Y^ line (13.2% vs 10.0%, respectively). No elevated survival was seen in the Cam3.II^ctrl^ line expressing only the silent shield mutations used at the guide RNA cut site.

### Southeast Asian K13 mutations do not impart a significant fitness impact on Dd2 parasites

Prior studies with isogenic gene-edited Southeast Asian lines have shown that certain K13 mutations can exert fitness costs, as demonstrated by reduced intra-erythrocytic asexual blood stage parasite growth (Straimer *et al.*, 2017; Nair *et al.*, 2018). To determine the fitness impact of the K13 mutations described above, we utilized an eGFP-based parasite competitive growth assay (Ross *et al.*, 2018). Dd2^E252Q^, Dd2^F446I^, Dd2^P553L^, Dd2^R561H^ or Dd2^P574L^ were co-cultured in 1:1 mixtures with an isogenic K13 WT eGFP^+^ Dd2 reporter line for 20 days (10 generations), and the proportion of eGFP^+^ parasites was assessed every two days. As controls, we included Dd2^WT^, Dd2^bsm^ and Dd2^C580Y^. These data provided evidence of a minimal impact with the F446I, P553L and C580Y mutations, with E252Q, R561H and P574L having greater fitness costs (**Figure 5C; Figure S4; Table S7**). Both C580Y and P553L displayed elevated RSA survival and minimal fitness cost in the Dd2 strain, providing optimal traits for dissemination (**Figure 5D**). We note that all fitness costs in Dd2 were considerably lower than those observed in our four African strains (**Figure 3**).

### Strain-dependent genetic background differences significantly RSA survival rates in culture-adapted Thai isolates

Given the earlier abundance of the R561H and E252Q alleles in border regions of Thailand and Myanmar, we next tested the impact of introducing these mutations into five Thai K13 WT isolates (Thai1-5). For comparison, we also edited C580Y into several isolates. These studies revealed a major contribution of the parasite genetic background in dictating the level of mutant K13-mediated ART resistance, as exemplified by the C580Y lines whose mean survival rates ranged from 2.1% to 15.4%. Trends observed for individual mutations were maintained across strains, with the R561H mutation consistently yielding moderate to high levels of *in vitro* resistance at or above the level of C580Y. Consistent with results for Dd2, introduction of E252Q did not result in significant increases in survival rates relative to isogenic K13 WT lines (**Figure 6A-E; Table S6**).

**Figure 6.**
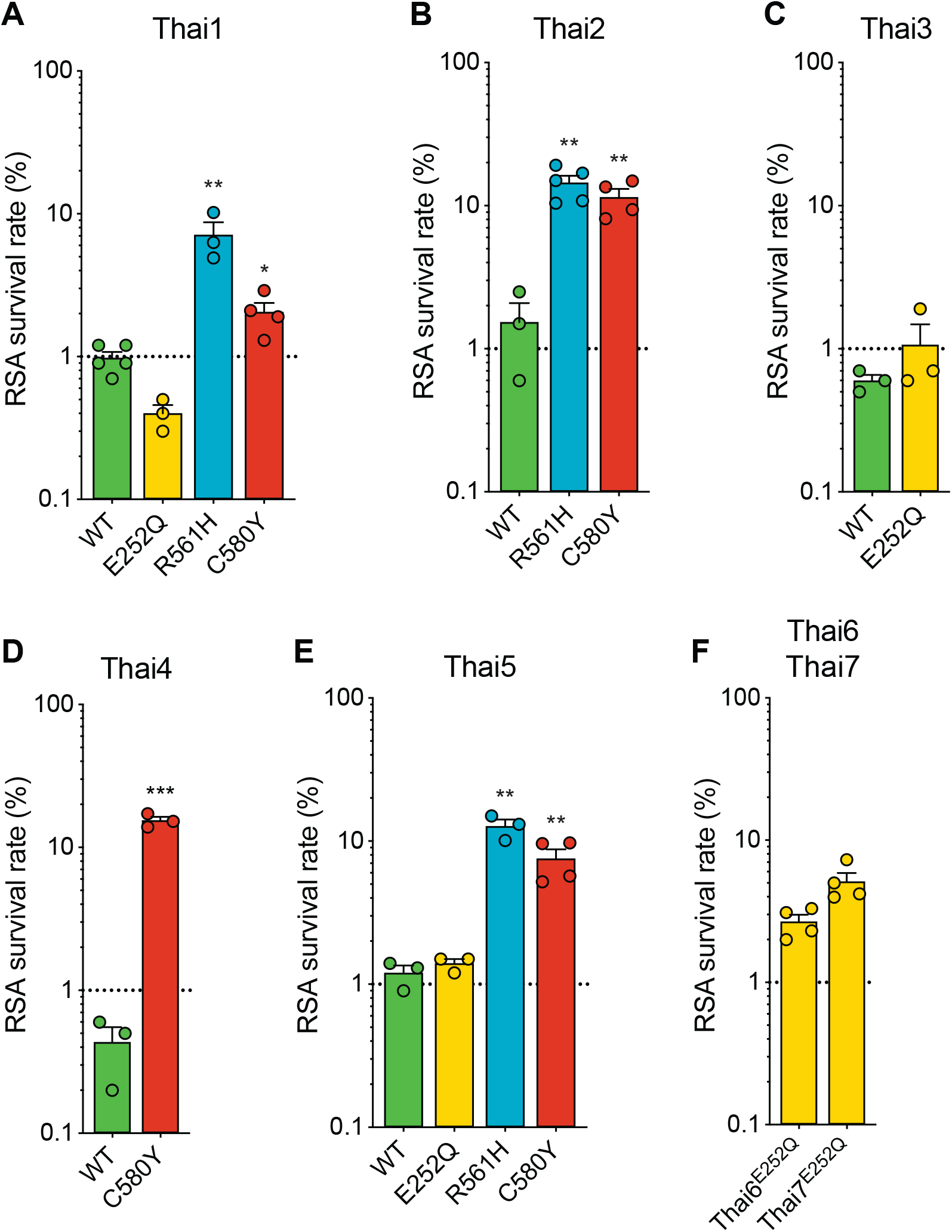
Thai isolates expressing mutant K13 display variable RSA survival rates. RSA survival rates for (A-E) *K13*-edited Thai isolates and (F) K13 E252Q unedited Thai lines, shown as means ± SEM (detailed in **Table S6**). Results were obtained from 3 to 7 independent experiments, each performed in duplicate. *P* values were determined by unpaired *t* tests and were calculated for mutant lines relative to the isogenic line expressing WT *K13*. * *P*<0.05; ** *P*<0.01; *** *P*<0.001.

We also profiled two unedited culture-adapted Thai isolates (Thai6^E252Q^ and Thai7^E252Q^) that express the K13 E252Q mutation, but that are otherwise K13 WT. Notably, both lines exhibited mean RSA survival rates significantly above the 1% threshold for ART sensitivity (2.7% for Thai6^E252Q^ and 5.1% for Thai7^E252Q^; **Figure 6F**). These data suggest that additional genetic factors present in these two Thai isolates are required for E252Q to manifest ART resistance.

### Mutations in the *P. falciparum* multidrug resistance protein 2 and ferredoxin genes do not modulate resistance to artemisinin *in vitro*

In a prior genome-wide association study of SE Asian parasites, K13-mediated ART resistance was associated with D193Y and T484I mutations in the ferredoxin (*fd*) and multidrug resistance protein 2 (*mdr2*) genes, respectively (Miotto *et al.*, 2015). To directly test the role of these mutations, we applied CRISPR/Cas9 editing to the Cambodian strains RF7^C580Y^ and Cam3.II^C580Y^, which both express K13 C580Y (**Figure S5**). Isogenic RF7 parasites expressing the mutant or wild-type fd residue at position 193 showed no change in RSA survival rates, either at 700 nM (averaging ~27%), or across a range of DHA concentrations down to 1.4 nM. Editing fd D193Y into K13 C580Y CamWT parasites (Straimer *et al.*, 2015) also had no impact on RSA survival (with mean RSA survival rates of 11-13%). Likewise, Cam3.II parasites maintained the same rate *of in vitro* RSA survival (mean 19-22%) irrespective of their *mdr2* allele. Silent shield mutations had no impact for either *fd* or *mdr2* (**Figure 7**; **Table S8**). These data suggest that the fd D193Y and mdr2 T484I mutations are markers of ART-resistant founder populations but themselves do not contribute directly to ART resistance.

**Figure 7.**
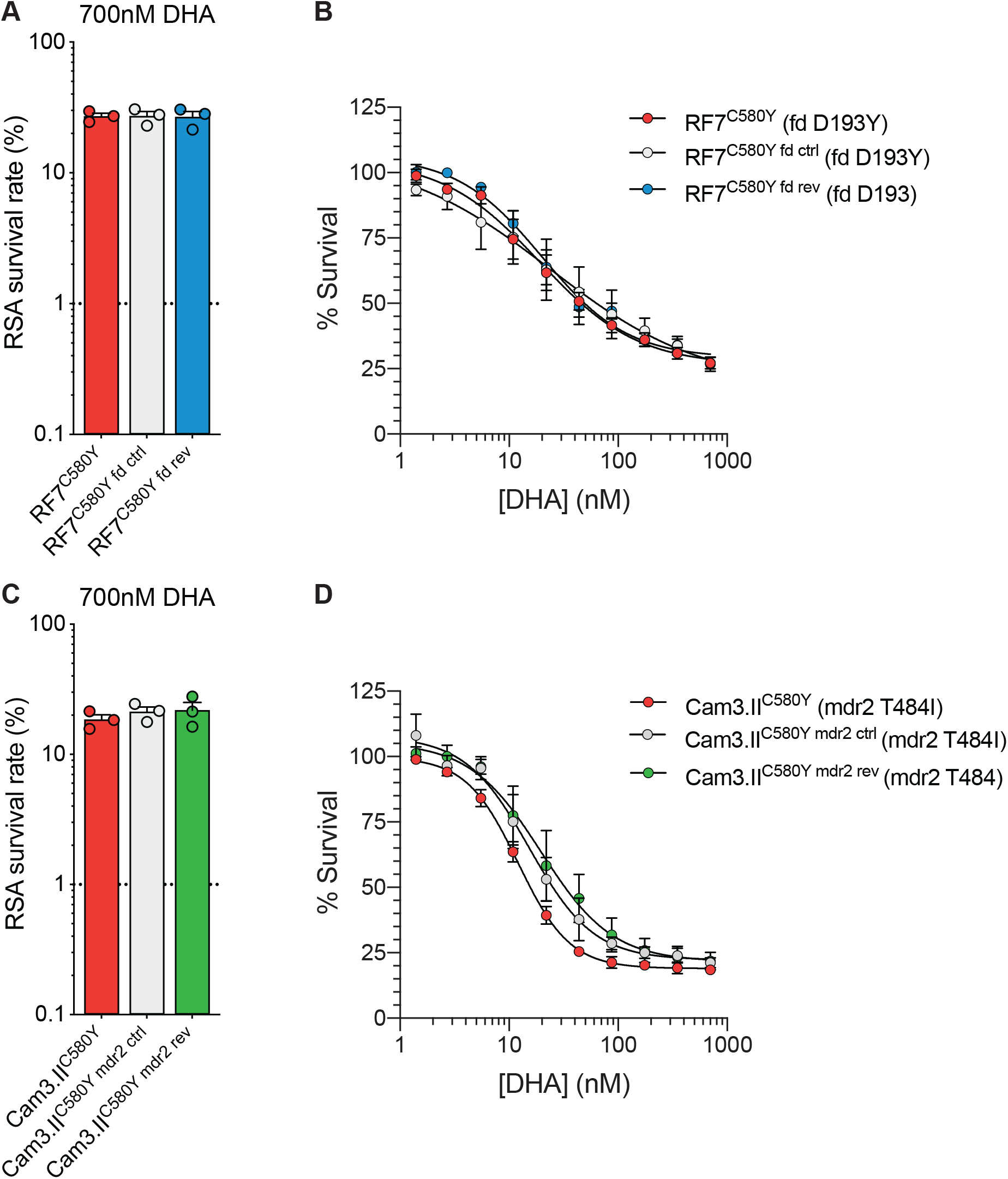
Ferrodoxin (*fd)* and multidrug resistance protein 2 (*mdr2)* mutations do not impact RSA survival in K13 C580Y parasites. (A, B) Results of RSA assays performed on the RF7^C580Y^ parental line that expresses the fd variant D193Y, a gene-edited control line with silent binding-site mutations at the *fd* gRNA cut site, and a gene-edited fd revertant D193 line. Gene-edited parasite lines were generated using CRISPR/Cas9. Early ring-stage parasites (0-3 hpi) were pulsed with serial dilutions of DHA (beginning with 700 nM DHA) for 6 h followed by drug washout. Results show (A) mean ± SEM survival at 700 nM, and (B) survival curves across the range of DHA concentrations. Survival rates were calculated relative to DMSO-treated parasites processed in parallel. Results were obtained from three independent experiments, each performed in duplicate. (C, D) RSAs performed on the Cam3.II^C580Y^ parental line (expressing the mdr2 variant T484I), a gene-edited control line with silent-binding site mutations at the *mdr2* gRNA cut site, and a gene-edited mdr2 revertant T484 line. Results shown in (C, D) were generated and presented as per (A, B). All values are provided in **Table S8**.

## Discussion

Mutant K13-mediated ART resistance has substantially compromised the efficacy of antimalarial treatments across SEA, and the relatively high prevalence of the R561H variant associated with delayed clearance in Rwanda now poses a threat to high-transmission settings in sub-Saharan Africa (Conrad and Rosenthal, 2019; Hanboonkunupakarn and White, 2020; Uwimana *et al.*, 2020; Bergmann *et al.*, 2021). Using gene editing and phenotypic analyses, we provide definitive evidence that the K13 R561H, M579I and C580Y mutations can confer *in vitro* ART resistance in several African strains. *In vitro* resistance, as defined using the RSA, was comparable between gene-edited African K13 R561H 3D7 parasites and Asian C580Y Dd2 and Cam3.II parasites. We also observed that K13 mutant African strains differed widely in their RSA survival rates. As an example, when introduced into the Tanzanian F32 and Ugandan UG815 strains, the C580Y mutation yielded 0.3% (not resistant) and 11.8% (highly resistant) RSA survival rates, respectively. These data suggest that F32 parasites lack additional genetic determinants that are required for mutant K13 to confer ART resistance. Nonetheless, our results provide conclusive evidence that multiple African strains present no core biological obstacle to becoming ART resistant upon acquiring K13 mutations.

Our spatio-temporal analysis of K13 sequence diversity in Cambodia highlights the emergence of C580Y in western Cambodia and its progressive replacement of other variants (Imwong *et al.*, 2020). The success of this mutation in SEA cannot be explained by resistance alone, as we previously reported that the less common R539T and I543T variants conferred greater ART resistance *in vitro* (Straimer *et al.*, 2015). Similarly, we now report that R561H and P553L yield equivalent degrees of ART resistance when compared with C580Y. Lower levels of resistance were observed with F446I and P574L, with the former predominating recently on the Thai-Myanmar border (Imwong *et al.*, 2020). In a recent study, F446I yielded no significant *in vitro* resistance in 3D7 parasites, although similar to our data this mutation was fitness-neutral (Siddiqui *et al.*, 2020). Of note, all four of these mutations, and others including C469Y, R622I and A675V, have now been detected in Africa and merit gene editing experiments in African strains [Warsame, 2019 #798;Kayiba, 2020 #796;Asua, 2020 #803].

Here we observed that the C580Y mutation exerts less of a fitness cost relative to other K13 variants, as measured in *K13*-edited Dd2 parasites co-cultured with an eGFP reporter line. These data suggest that C580Y might be favored in part because of a less detrimental impact on asexual blood stage growth rates. The most detrimental impact on growth was observed with E252Q, which earlier predominated near the Thailand-Myanmar border but was later overtaken by C580Y, as well as R561H, which progressively disappeared over time in SEA (Phyo *et al.*, 2016). In our study C580Y produced an optimal combination of no measurable fitness cost and relatively high RSA survival rates in Dd2 parasites. R561H, however, showed slightly improved fitness relative to C580Y in paired isogenic parasites from Thailand (the NHP4302 strain) (Nair *et al.*, 2018), providing evidence that both fitness and resistance are strain-dependent.

Consistent with these findings, we observed substantial fitness costs with the K13 C580Y mutation in four African strains. The largest growth defect was observed with the edited UG815 C580Y line that also yielded the highest level of ART resistance. These data suggest that K13 C580Y may not easily take hold in Africa where, unlike in SEA, infections are often highly polyclonal, generating intra-host competition that impacts a strain’s ability to succeed at the population level. In addition, individuals in highly-endemic African settings generally have high levels of acquired immunity, potentially preventing infection by relatively unfit parasites, and often have asymptomatic infections that go untreated and are thus less subject to selective drug pressure, compared with individuals in SEA. This situation recalls the history of chloroquine use in Africa, where fitness costs caused by mutations in the primary resistance determinant PfCRT resulted in the rapid resurgence of wild-type parasites following the implementation of other first-line antimalarial therapies (Kublin *et al.*, 2003; Laufer *et al.*, 2006; Ord *et al.*, 2007; Frosch *et al.*, 2014). It remains to be determined whether mutations such as R561H, emerging in Rwanda, can ameliorate the fitness cost observed with other K13 variants in African strains.

Further research is also required to define secondary genetic determinants that could augment mutant K13-mediated ART resistance and to explore other potential mediators of resistance. The latter include mutations in AP-2μ, UBP-1 and Pfcoronin, which can modulate *P. falciparum* ART susceptibility *in vitro* and merit further investigation (Demas *et al.*, 2018; Henrici *et al.*, 2019; Sutherland *et al.*, 2020). Data provided herein argue against a direct role for mutations in *fd* and *mdr2*, earlier associated with mutant K13-mediated resistance in SEA (Miotto *et al.*, 2015). We note that *P. falciparum* population structures in Africa tend to be far more diverse than in the epicenter of resistance in Cambodia, where parasite strains are highly sub-structured into a few lineages that can readily maintain complex genetic traits (Amato *et al.*, 2018). A requirement to transmit mutant K13 and additional determinants of resistance in African malaria-endemic settings, where genetic outcrossing is the norm, would predict that ART resistance will spread more gradually than in SEA.

Another impediment to the dissemination of ART resistance in Africa is the continued potent activity of lumefantrine, the partner drug in the first line treatment artemether-lumefantrine. This situation contrasts with SEA where ART-resistant parasites also developed high-level resistance to the partner drug PPQ, with widespread treatment failures enabling the dissemination of multidrug-resistant strains (Conrad and Rosenthal, 2019; van der Pluijm *et al.*, 2019). These data call for continent-wide monitoring for the emergence and spread of mutant K13 in Africa, and for studies of whether its emergence in Rwanda is a harbinger of subsequent partner drug resistance and ACT treatment failure.

## Materials and Methods

### *P. falciparum* parasite *in vitro* culture

*Plasmodium falciparum* asexual blood-stage parasites were cultured in human erythrocytes at 3% hematocrit in RPMI-1640 medium supplemented with 2 mM L-glutamine, 50 mg/L hypoxanthine, 25 mM HEPES, 0.225% NaHCO3, 10 mg/L gentamycin and 0.5% w/v Albumax II (Invitrogen). Parasites were maintained at 37°C in 5% O_2_, 5% CO_2_, and 90% N2. Cultures were monitored by light microscopy of methanol-fixed, Giemsa-stained blood smears. The geographic origin and year of culture adaptation for lines employed herein are described in **Table 1** and **Table S2**.

### Whole-genome sequencing of parental lines

To define the genome sequences of our *P. falciparum* lines used for transfection, we lysed parasites in 0.05% saponin, washed them with 1×PBS, and purified genomic DNA (gDNA) using the QIAamp DNA Blood Midi Kit (Qiagen). DNA concentrations were quantified by NanoDrop (Thermo Scientific) and Qubit (Invitrogen) prior to sequencing. 200 ng of gDNA was used to prepare sequencing libraries using the Illumina Nextera DNA Flex library prep kit with dual indices. Samples were multiplexed and sequenced on an Illumina MiSeq to obtain 300 bp paired-end reads at an average of 50× depth of coverage. Sequence reads were aligned to the *P. falciparum* 3D7 reference genome (PlasmoDB version 36) using Burrow-Wheeler Alignment. PCR duplicates and unmapped reads were filtered out using Samtools and Picard. Reads were realigned around indels using GATK RealignerTargetCreator and base quality scores were recalibrated using GATK BaseRecalibrator. GATK HaplotypeCaller (version 3.8) was used to identify all single nucleotide polymorphisms (SNPs). These SNPs were filtered based on quality scores (variant quality as function of depth QD > 1.5, mapping quality > 40, min base quality score > 18) and read depth (> 5) to obtain high-quality SNPs, which were annotated using snpEFF. Integrated Genome Viewer was used to visually verify the presence of SNPs. BIC-Seq was used to check for copy number variations using the Bayesian statistical model (Xi *et al.*, 2011). Copy number variations in highly polymorphic surface antigens and multi-gene families were removed as these are prone to copy number changes with *in vitro* culture.

These whole-genome sequencing data were used to determine the genotypes of the antimalarial drug resistance loci *pfcrt*, *mdr1*, *dhfr* and *dhps* (Haldar *et al.*, 2018). We also genotyped *fd*, *arps10*, *mdr2*, *ubp1,* and *ap-2μ*, which were previously associated with ART resistance (Henriques *et al.*, 2014; Miotto *et al.*, 2015; Cerqueira *et al.*, 2017; Adams *et al.*, 2018). These results are described in **Table S2**.

### Cloning of *K13, fd* and *mdr2* plasmids

Zinc-finger nuclease-meditated editing of select mutations in the *K13* locus (**Table S3**; **Table S6**) was performed as previously described (Straimer *et al.*, 2015). CRISPR/Cas9 editing of *K13* mutations was achieved using the pDC2-cam-coSpCas9-U6-gRNA-h*dhfr* all-in-one plasmid that contains a *P. falciparum* codon-optimized Cas9 sequence, a human dihydrofolate reductase (h*dhfr*) gene expression cassette (conferring resistance to WR99210) and restriction enzyme insertion sites for the guide RNA (gRNA) and donor template (White *et al.*, 2019). A K13 propeller domain-specific guide gRNA was introduced into this vector at the BbsI restriction sites using the oligo pair p1+p2 (**Table S9**) using T4 DNA ligase (New England BioLabs). Oligos were phosphorylated and annealed prior to cloning. A *K13* donor template consisting of a 1.5 kb region of the *K13* coding region including the entire propeller domain was amplified using the primer pair p3+p4 (**Table S9**) and cloned into the pGEM T-easy vector system (Promega). This donor sequence was subjected to site-directed mutagenesis in the pGEM vector to introduce silent binding-site mutations at the Cas9 cleavage site using the primer pair p5+p6, and to introduce allele-specific mutations using the primer pairs (p7 to p20) described in **Table S9**. *K13* donor sequences were amplified from the pGEM vector using the primer pair p21+p22 and sub-cloned into the pDC2-cam-coSpCas9-U6-gRNA-h*dhfr* plasmid at the EcoRI and AatII restriction sites by In-Fusion® Cloning (Takara). The final plasmids were then sequenced using primers p23 to p25 (**Table S9**). A schematic showing the method of *K13* plasmid construction can be found in **Figure S1**.

CRISPR/Cas9 editing of *fd* and *mdr2* was performed using a separate all-in-one plasmid, pDC2-cam-Cas9-U6-gRNA-h*dhfr*, generated prior to the development of the codon-optimized version used above for *K13* (Lim *et al.*, 2016). Cloning was performed as for *K13*, except for gRNA cloning that was performed using In-Fusion® cloning (Takara) rather than T4 ligase. Cloning of gRNAs was performed using primer pair p29/p30 for *fd* and p42/p43 for *mdr2*. Donor templates were amplified and cloned into the final vector using the primer pairs p31/p32 for *fd* and p44+p45 for *mdr2*. Site-directed mutagenesis was performed using the allele-specific primer pairs p33+p34 or p35+p36 for *fd*, and p46+p47 or p48+p49 for *mdr2*. All final plasmids (both *fd*- and *mdr2*-specific) were sequenced using the primer pair p37+p38 (**Table S9**; **Table S10**). Schematic representations of final plasmids are shown in **Figure S5**.

### Generation of *K13*, *fd* and *mdr2* gene-edited parasite lines

Gene-edited lines were generated by electroporating ring-stage parasites at 5-10% parasitemia with 50 μg of purified circular plasmid DNA resuspended in Cytomix. Transfected parasites were selected by culturing in the presence of WR99210 (Jacobus Pharmaceuticals) for six days post electroporation. Parental lines harboring 2-3 mutations in the *P. falciparum* dihydrofolate reductase (*dhfr*) gene were exposed to 2.5 nM WR99210, while parasites harboring four *dhfr* mutations were selected under 10 nM WR99210 (see **Table S2**). Parasite cultures were monitored for recrudescence by microscopy for up six weeks post electroporation. To test for successful editing, the *K13* locus was amplified directly from whole blood using the primer pair p26+p27 (**Table S9**) and the MyTaq™ Blood-PCR Kit (Bioline). Primer pairs p39+p40 and p50+p51 were used to amplify *fd* and *mdr2*, respectively (**Table S9**). PCR products were submitted for Sanger sequencing using the PCR primers as well as primer p28 in the case of *K13*, p41 (*fd*) or p52 (*mdr2*) (**Table S9**). Bulk-transfected cultures showing evidence of editing by Sanger sequencing were cloned by limiting dilution.

### Parasite synchronization, ring-stage survival assays (RSAs) and flow cytometry

Synchronized parasite cultures were obtained by exposing predominantly ring-stage cultures to 5% D-Sorbitol (Sigma) for 15 min at 37°C to remove mature parasites. After 36 h of subsequent culture, multinucleated schizonts were either purified over a density gradient consisting of 75% Percoll (Sigma). Purified schizonts were incubated with fresh RBCs for 3h, and early rings (0-3 hours post invasion; hpi) were treated with 5% D-Sorbitol to remove remaining schizonts.

*In vitro* RSAs were conducted as previously described, with minor adaptations (Straimer *et al.*, 2015). Briefly, tightly synchronized 0-3 hpi rings were exposed to a pharmacologically-relevant dose of 700 nM DHA or 0.1% dimethyl sulfoxide (DMSO; vehicle control) for 6 h at 1% parasitemia and 2% hematocrit, washed three times with RPMI medium to remove drug, transferred, and cultured for an additional 66 h in drug-free medium. Removal of media and resuspension of parasite cultures was performed on a Freedom Evo 100 liquid-handling instrument (Tecan). Parasitemias were measured at 72 h by flow cytometry (see below) with at least 50,000 events captured per sample. Parasite survival was expressed as the percentage value of the parasitemia in DHA-treated samples divided by the parasitemia in DMSO-treated samples processed in parallel. We considered any RSA mean survival rates <2% to be ART sensitive.

Flow cytometry was performed on an BD Accuri^TM^ C6 Plus cytometer with a HyperCyt plate sampling attachment (IntelliCyt), or on an iQue3® Screener Plus cytometer (Sartorius). Cells were stained with 1×SYBR Green (Invitrogen) and 100 nM MitoTracker DeepRed (Invitrogen) for 30 min and diluted in 1×PBS prior to sampling. Percent parasitemia was determined as the percentage of MitoTracker^-^positive and SYBR Green-positive cells. For RSAs, >50,000 events were captured per well.

### TaqMan allelic discrimination real-time (quantitative) PCR-based fitness assays

Fitness assays with African *K13*-edited parasite lines were performed by co-culturing isogenic wild-type unedited and mutant edited parasites in 1:1 ratios. Assays were initiated with tightly synchronized trophozoites. Final culture volumes were 3 mL. Cultures were maintained in 12-well plates and monitored every four days over a period of 40 days (20 generations) by harvesting at each time point a fraction of each co-culture for saponin lysis. gDNA was then extracted using the QIAamp DNA Blood Mini Kit (Qiagen). The percentage of the WT or mutant allele in each sample was determined in TaqMan allelic discrimination real-time PCR assays. TaqMan primers (forward and reverse) and TaqMan fluorescence-labeled minor groove binder probes (FAM or HEX, Eurofins) are described in **Table S11**. Probes were designed to specifically detect the K13 M579I or C580Y propeller mutations. The efficiency and sensitivity of the TaqMan primers was assessed using standard curves comprising 10-fold serially diluted templates ranging from 10 ng to 0.001 ng. Robustness was demonstrated by high efficiency (88-95%) and R^2^ values (0.98-1.00). The quantitative accuracy in genotype calling was assessed by performing multiplex qPCR assays using mixtures of WT and mutant plasmids in fixed ratios (0:100, 20:80, 40:60, 50:50, 60:40, 80:20, 100:0). Triplicate data points clustered tightly, indicating high reproducibility in the data across the fitted curve (R^2^ = 0.89 to 0.91).

Purified gDNA from fitness co-cultures was subsequently amplified and labeled using the primers and probes described in **Table S11**. qPCR reactions for each sample were run in triplicate. 20 μL reactions consisted of 1×QuantiFAST reaction mix containing ROX reference dye (Qiagen, Germany), 0.66 µM of forward and reverse primers, 0.16 µM FAM-MGB and HEX-MGB TaqMan probes, and 10 ng genomic DNA. Amplification and detection of fluorescence were carried out on the QuantStudio 3 (Applied Biosystems) using the genotyping assay mode. Cycling conditions were as follows: 30s at 60°C; 5 min at 95°C; and 40 cycles of 30s at 95°C and 1 min at 60°C for primer annealing and extension. Every assay was run with positive controls (WT or mutant plasmids at different fixed ratios). No-template negative controls (water) in triplicates were processed in parallel. Rn, the fluorescence of the FAM or HEX probe, was normalized to the fluorescence signal of the ROX reporter dye. Background-normalized fluorescence (Rn minus baseline, or ∆Rn) was calculated as a function of cycle number.

To determine the WT or mutant allele frequency in each sample, we first confirmed the presence of the allele by only retaining values where the threshold cycle (Ct) of the sample was less than the no-template control by at least three cycles. Next, we subtracted the ∆Rn of the samples from the background ∆Rn of the no-template negative control. We subsequently normalized the fluorescence to 100% using the positive control plasmids to obtain the percentage of the WT and mutant alleles for each sample. The final percentage of the mutant allele was defined as the average of these two values: the normalized percentage of the mutant allele, and 100% minus the normalized percentage of the wild-type allele.

### eGFP-based fitness assays

Fitness assays with Dd2 parasite lines were performed as previously described (Ross *et al.*, 2018). Briefly, *K13*-edited parasite lines were co-cultured in 1:1 ratios with an eGFP-positive (eGFP^+^) Dd2 reporter line. Fitness assays were initiated with tightly synchronized trophozoites in 96-well plates, with 200 μL culture volumes. Percentages of eGFP^+^ parasites were monitored by flow cytometry every two days over a period of 20 days (10 generations). Flow cytometry was performed as written above, except that only 100 nM MitoTracker DeepRed staining was used to detect total parasitemias, since SYBR Green and eGFP fluoresce in the same channel.

### Fitness Costs

The fitness cost associated with a line expressing a given K13 mutation was calculated was calculated relative to its isogenic WT counterpart using the following equation:

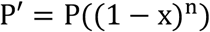

where P’ is equal to the parasitemia at the assay endpoint, P is equal to the parasitemia on day 0, n is equal to the number of generations from the assay start to finish, and x is equal to the fitness cost. This equation assumes 100% growth for the WT comparator line. For qPCR and GFP-based fitness assays, days 32 and 20 were set as the assay endpoints, resulting in the number of parasite generations (n) being set to 16 and 10, respectively.

## Acknowledgments

We thank Dr. Pascal Ringwald (World Health Organization) for his support and feedback. DAF gratefully acknowledges the US National Institutes of Health (R01 AI109023), the Department of Defense (W81XWH1910086) and the Bill & Melinda Gates Foundation (OPP1201387) for their financial support. BHS was funded in part by T32 AI106711 (PD: D. Fidock). SM is a recipient of a Human Frontiers of Science Program Long-Term Fellowship. CHC was supported in part by the NIH (R01 AI121558; PI: Jonathan Juliano). FN is supported by the Wellcome Trust of Great Britain (Grant ID: 106698). TJCA acknowledges funding support from the NIH (R37 AI048071). DAF and DM gratefully acknowledge the World Health Organization for their funding. We thank the following individuals for their kind help with the *K13*-genotyped samples – Chad: Ali S. Djiddi, Mahamat S. I. Diar, Kodbessé Boulotigam, Mbanga Djimadoum, Hamit M. Alio, Mahamat M. H. Taisso, Issa A. Haggar; Burkina Faso: TES 2017-2018 team and the US President’s Malaria Initiative through the Improving Malaria Care Project as the funding agency for the study in Burkina Faso, Chris-Boris G. Panté-Wockama; Burundi: Dismas Baza; Tanzania: Mwaka Kakolwa, Celine Mandara, Tanzania TES coordination team for the Ministry of Health; Sierra Leone: Anitta R. Y. Kamara, Foday Sahr, Mohamed Samai; The Gambia: Balla Kandeh, Joseph Okebe, Serign J. Ceesay, Baboucarr Babou, Emily Jagne, Alsan Jobe; Congo: Brice S. Pembet, Jean M. Youndouka; Somalia: Jamal Ghilan Hefzullah Amran, Abdillahi Mohamed Hassan, Abdikarim Hussein Hassan and Ali Abdulrahman; Rwanda: extended TES team for the Malaria and Other Parasitic Diseases Division, Rwanda Biomedical Centre.

