## Supplemental Figures and Tables for "*Plasmodium falciparum* K13 mutations in Africa and Asia present varying degrees of artemisinin resistance and an elevated fitness cost in African parasites"

#### Table of Contents

**Figure S1.** CRISPR/Cas9 strategy for editing the K13 locus………………………...page 3

**Figure S2.** Crystal structure of K13 propeller domain showing positions of mutated residues………………………………………………………………………………page 4

**Figure S3.** Regions of sample collection in Cambodia for *K13* sequencing……….page 5

**Figure S4.** Southeast Asian K13 mutations result in minor *in vitro* growth defects in Dd2 parasites, with the exception of the C580Y and P553L mutations…………………..page 6

**Figure S5.** CRISPR/Cas9 strategies for editing the ferrodoxin (fd) and multidrug resistance protein 2 (mdr2) loci……………………………………………………….page 7

**Table S1.** Distribution of *K13* alleles over time in African countries.………………...page 8

**Table S2.** Geographic origin and drug resistance genotypes of *Plasmodium falciparum* clinical isolates and reference lines employed in this study…………………………..page 9

**Table S3.** Ring-stage survival (RSA) assay data for edited parasites and controls (African strains)……………………………………………………………………………...page 10

### Figure S1

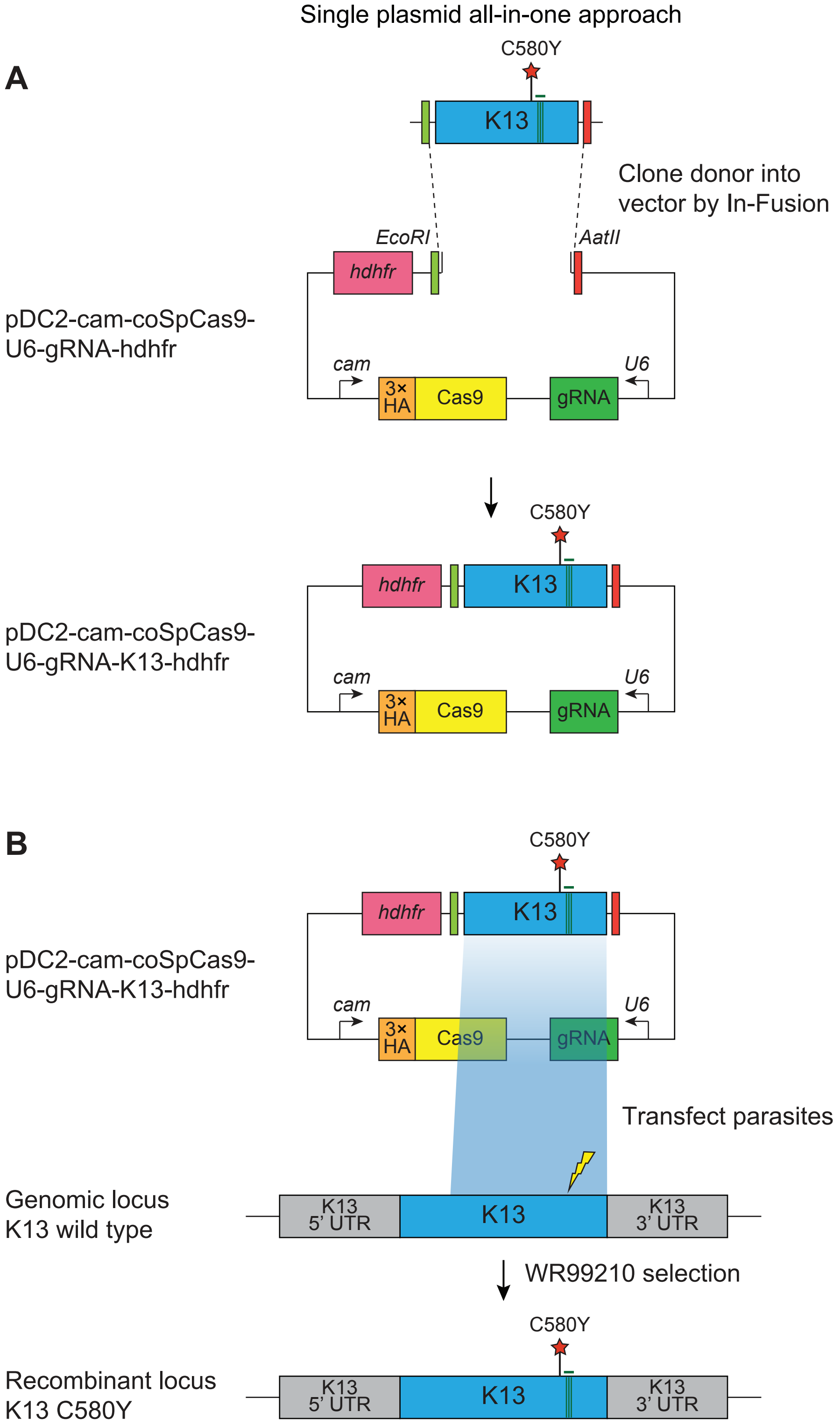

**Figure S1. CRISPR/Cas9 strategy for editing the K13 locus.**
All-in-one plasmid approach used for CRISPR/Cas9-mediated K13 gene editing, consisting of a K13-specific donor template for homology-directed repair, a K13-specific gRNA expressed from the U6 promoter, a Cas9 cassette with expression driven by the calmodulin (cam) promoter, and a selectable marker (human dhfr, conferring resistance to the antimalarial WR99210 that inhibits P. falciparum dhfr). The Cas9 sequence was codon-optimized for improved expression in P. falciparum. Donors coding for specific mutations of interest (e.g. K13 C580Y, red star) were generated by site-directed mutagenesis (SDM) of the K13 wild-type donor sequence. Green bars indicate the presence of silent binding-site mutations that were introduced by SDM to protect the edited locus from further cleavage. The lightning bolt indicates the location of the cut site in the genomic target locus. Primers used for SDM and cloning and transfection plasmids are described in Table S9 and Table S10, respectively.

### Figure S2

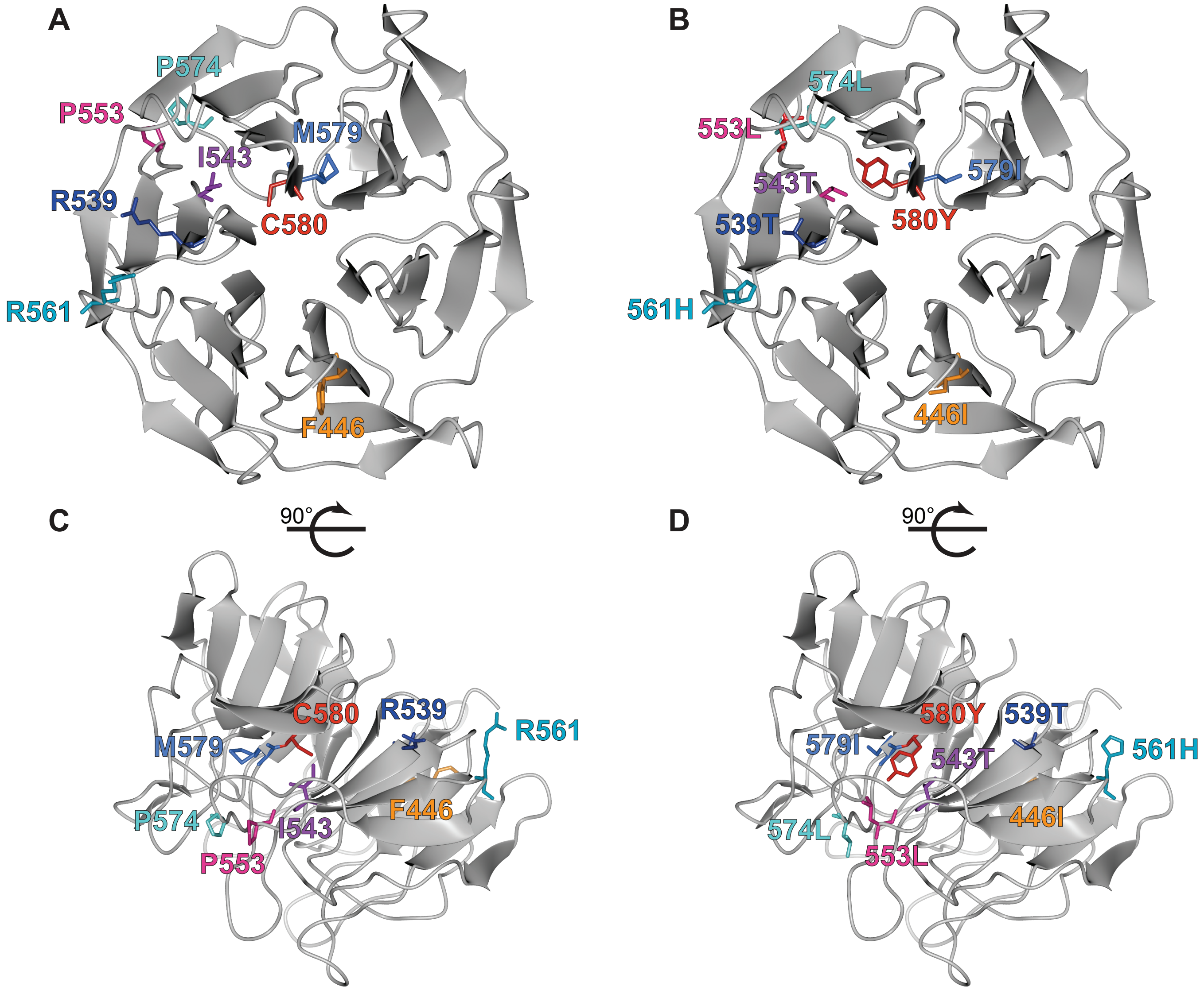

**Figure S2. Crystal structure of K13 propeller domain showing positions of mutated residues.** (A, B) Top and (C, D) side views of the crystal structure of the K13 propeller domain (PDB ID: 4YY8), highlighting residues of interest (F446I, orange; R539T, dark blue; I543T, purple; P553L, pink; R561H, dark turquoise; P574L, light turquoise; M579I medium blue; C580Y, red). Structures shown in (A) and (C) show wild-type residues while (B) and (D) show mutated residues.

### Figure S3

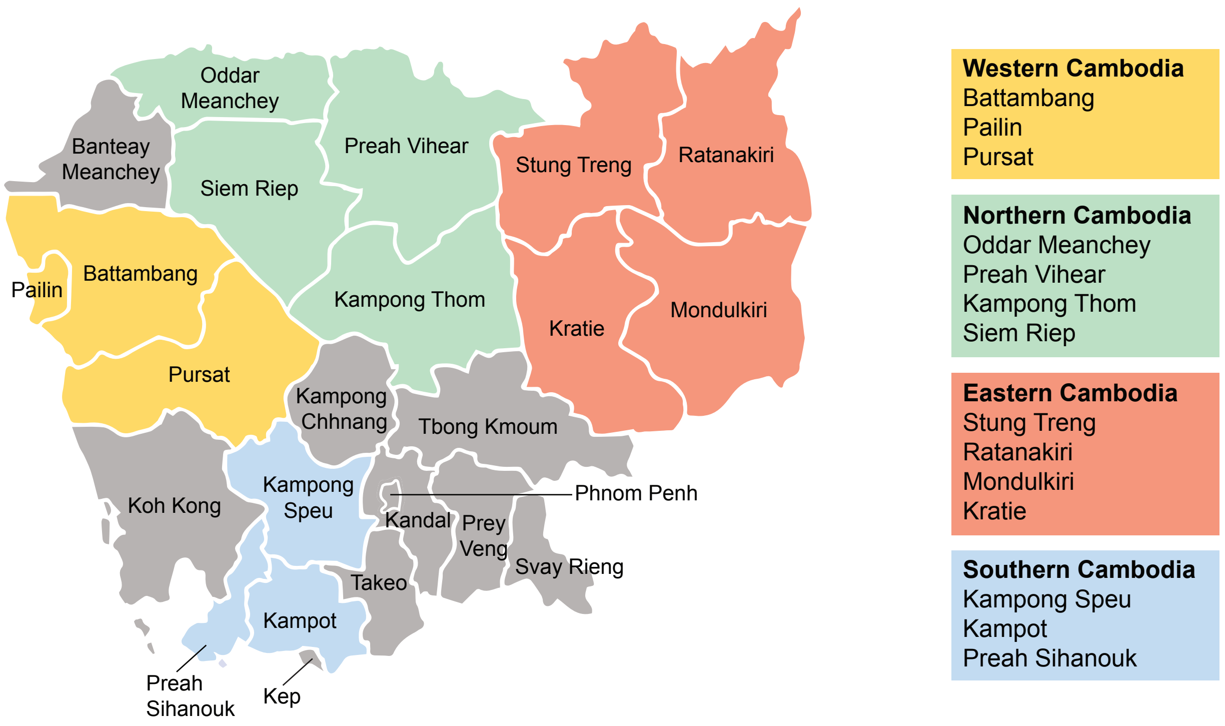

**Figure S3. Regions of sample collection in Cambodia for K13 sequencing.**
Map depicting the four regions of Cambodia (Western, Northern, Eastern, and Southern) in which samples were collected between 2001 and 2017 for K13 genotyping. Genotyping data are presented in Figure 4 and are tabulated in Table S5.

### Figure S4

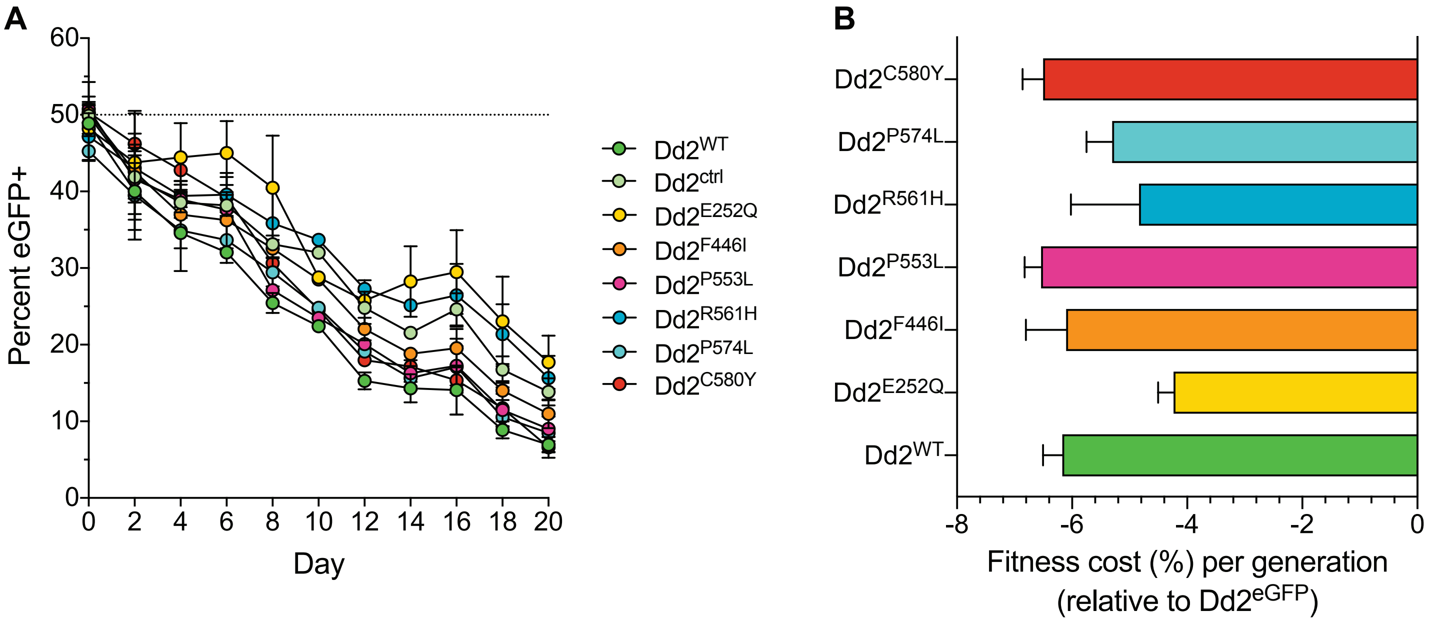

**Figure S4. Southeast Asian K13 mutations result in minor in vitro growth defects in Dd2 parasites, with the exception of the C580Y and P553L mutations.**
(A) Percentage of eGFP+ parasites over time in parasite cultures in which an eGFP-expressing Dd2 line was co-cultured in a 1:1 mixture with either the Dd2 K13 WT parental line ($Dd2^{WT}$) or individual Dd2 gene-edited K13 mutant lines. Parasite lines were co-cultured over a period of 20 days and percentages of eGFP+ parasites were determined by flow cytometry. Data are shown as means ± SEM (detailed in Table S7). Results were obtained from three independent experiments, each performed in triplicate. (B) Percent reduction in growth rate per 48 h generation, termed the fitness cost, are shown as mean ± SEM values for each mutant line relative to the $Dd2^{eGFP}$ line.

#### Figure S5

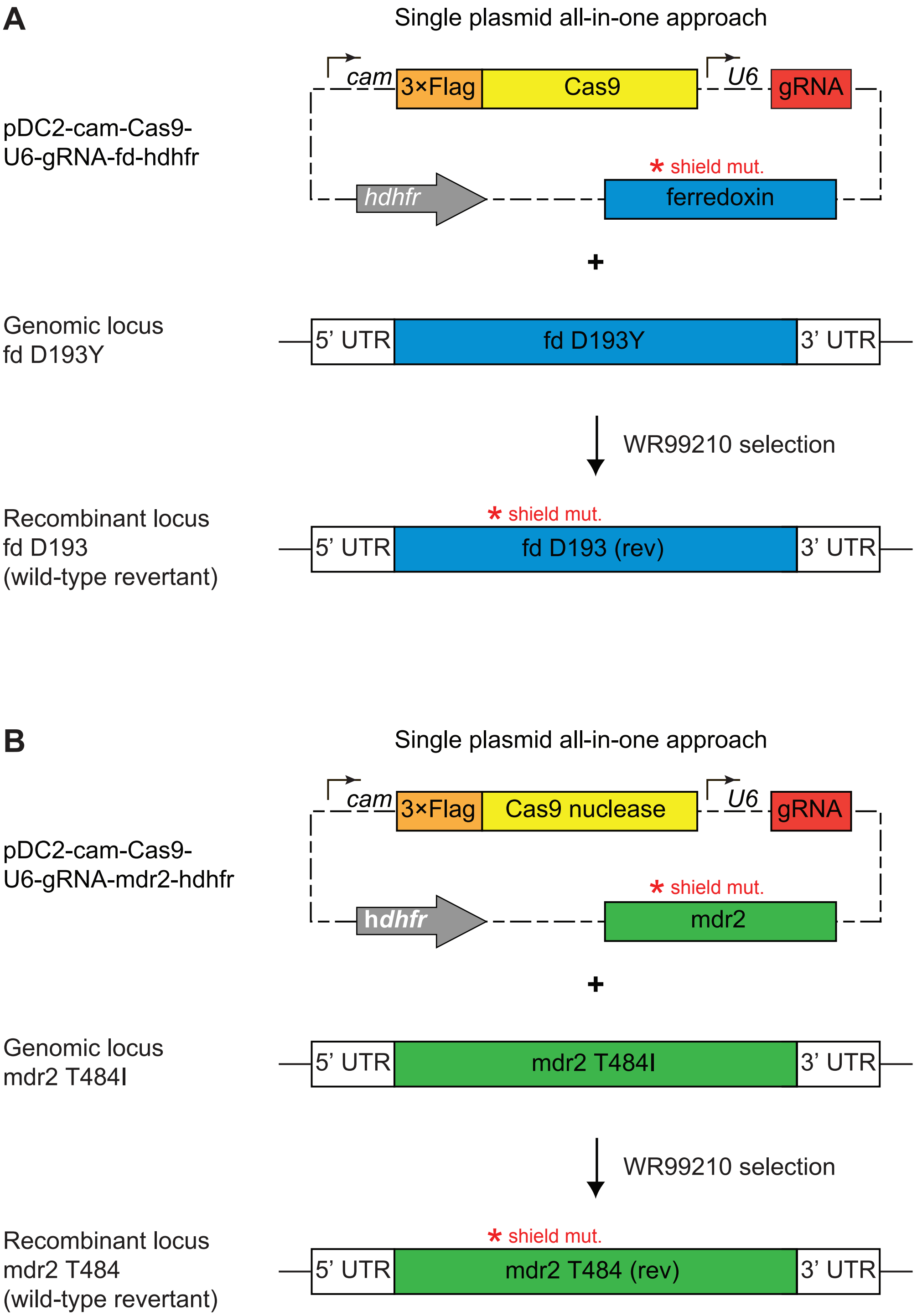

**Figure S5. CRISPR/Cas9 strategy for editing the ferrodoxin (fd) and multidrug resistance protein 2 (mdr2) loci.**
All-in-one plasmid approaches used for CRISPR/Cas9-mediated editing of (A) the ferredoxin (fd) locus or (B) the multidrug resistance protein 2 (mdr2) locus. Plasmids consisted of a fd (A) or mdr2 (B) specific donor template for homology-directed repair, a locus-specific gRNA expressed from the U6 promoter, a Cas9 cassette with expression driven by the cam promoter, and a selectable marker (hdhfr, conferring resistance to WR99210). Donors coding for specific mutations of interest (i.e. fd D193Y or mdr2 T484I) were generated by site-directed mutagenesis (SDM) of the WT donor sequences. Red stars indicate the presence of silent binding-site mutations (also introduced by SDM) used to protect edited loci from further cleavage. Primers used for SDM and cloning and transfection plasmids are described in Table S9 and Table S10, respectively.

**Table S1. Distribution of *K13* alleles over time in African countries.**

| Year(s) of sample collection | 2014, 2016 | 2016 | 2017-2018 | 2015 | 2017-2019 | 2014, 2015, 2017, 2019 | 2014 | 2015, 2016, 2019 | 2011, 2012, 2015 | 2012-2015 | 2016-2017 | |
|---|---|---|---|---|---|---|---|---|---|---|---|---|
| K13 | The Gambia | Sierra Leone[a] | Burkina Faso | Chad[b] | CAR[c] | Rep. of the Congo | Equatorial Guinea[d] | Burundi | Tanzania[e] | Rwanda[f] | Somalia[g] | Total |
| WT | 308 | 270 | 352 | 30 | 183 | 316 | 96 | 382 | 220 | 871 | 137 | 3165 |
| SYN | 3 | 2 | 8 | 1 | 3 | 12 | 0 | 7 | 8 | 11 | 0 | 55 |
| M460I | 0 | 0 | 0 | 0 | 0 | 0 | 0 | 0 | 0 | 1 | 0 | 1 |
| L463S | 0 | 0 | 0 | 0 | 0 | 0 | 0 | 0 | 1 | 0 | 0 | 1 |
| C469F | 0 | 0 | 0 | 0 | 0 | 0 | 0 | 0 | 0 | 1 | 0 | 1 |
| C469Y | 0 | 0 | 0 | 0 | 0 | 0 | 0 | 0 | 0 | 1 | 0 | 1 |
| M476I | 0 | 0 | 0 | 0 | 0 | 0 | 0 | 0 | 1 | 0 | 0 | 1 |
| V487I | 0 | 0 | 0 | 0 | 0 | 0 | 0 | 0 | 0 | 1 | 0 | 1 |
| G496S | 0 | 0 | 0 | 0 | 0 | 0 | 0 | 0 | 1 | 0 | 0 | 1 |
| V510M | 0 | 0 | 0 | 0 | 0 | 0 | 0 | 0 | 1 | 0 | 0 | 1 |
| R513L | 0 | 0 | 0 | 0 | 0 | 0 | 0 | 0 | 0 | 1 | 0 | 1 |
| V517I | 0 | 0 | 2 | 0 | 0 | 0 | 0 | 0 | 0 | 0 | 0 | 2 |
| V555A | 0 | 0 | 0 | 0 | 0 | 0 | 0 | 0 | 0 | 6 | 0 | 6 |
| E556K | 0 | 0 | 0 | 0 | 0 | 0 | 0 | 0 | 1 | 0 | 0 | 1 |
| A557S | 0 | 0 | 0 | 0 | 0 | 5 | 0 | 0 | 0 | 0 | 0 | 5 |
| R561H | 0 | 0 | 0 | 0 | 0 | 0 | 0 | 0 | 0 | 20 | 0 | 20 |
| M562T | 0 | 0 | 0 | 0 | 0 | 0 | 0 | 0 | 1 | 0 | 0 | 1 |
| P574L | 0 | 0 | 0 | 0 | 0 | 0 | 0 | 0 | 0 | 1 | 0 | 1 |
| R575I | 0 | 0 | 0 | 0 | 0 | 0 | 0 | 0 | 0 | 2 | 0 | 2 |
| A578S | 0 | 3 | 3 | 0 | 0 | 0 | 2 | 0 | 0 | 2 | 0 | 10 |
| A578V | 0 | 0 | 0 | 0 | 0 | 0 | 0 | 0 | 0 | 1 | 0 | 1 |
| V589A | 0 | 0 | 1 | 0 | 0 | 0 | 0 | 0 | 0 | 0 | 0 | 1 |
| E602D | 0 | 0 | 0 | 0 | 0 | 0 | 0 | 0 | 1 | 0 | 0 | 1 |
| V603I | 0 | 0 | 0 | 0 | 0 | 1 | 0 | 0 | 0 | 0 | 0 | 1 |
| E605K | 0 | 0 | 0 | 0 | 0 | 0 | 0 | 0 | 0 | 1 | 0 | 1 |
| R622I | 0 | 0 | 0 | 0 | 0 | 0 | 0 | 0 | 0 | 0 | 1 | 1 |
| A626E | 0 | 0 | 0 | 0 | 0 | 0 | 0 | 0 | 0 | 1 | 0 | 1 |
| N629Y | 0 | 0 | 1 | 0 | 0 | 0 | 0 | 0 | 0 | 0 | 0 | 1 |
| H644L | 1 | 0 | 0 | 0 | 0 | 0 | 0 | 0 | 0 | 0 | 0 | 1 |
| I646T | 0 | 3 | 0 | 0 | 0 | 0 | 0 | 0 | 0 | 0 | 0 | 3 |
| E651K | 0 | 0 | 0 | 0 | 0 | 0 | 0 | 0 | 0 | 1 | 0 | 1 |
| Y653N | 0 | 0 | 0 | 0 | 1 | 0 | 0 | 0 | 0 | 0 | 0 | 1 |
| V666I | 1 | 0 | 0 | 0 | 0 | 0 | 0 | 0 | 0 | 0 | 0 | 1 |
| P667R | 0 | 0 | 0 | 0 | 0 | 0 | 0 | 0 | 0 | 4 | 0 | 4 |
| V520I+V637I | 0 | 0 | 0 | 0 | 0 | 1 | 0 | 0 | 0 | 0 | 0 | 1 |
| V568M+V603I | 0 | 0 | 0 | 0 | 0 | 1 | 0 | 0 | 0 | 0 | 0 | 1 |
| G592E+V637I | 0 | 0 | 0 | 0 | 0 | 0 | 0 | 0 | 0 | 1 | 0 | 1 |
| Total | 313 | 278 | 367 | 31 | 187 | 336 | 98 | 389 | 235 | 927 | 138 | 3299 |
| % WT[h] | 99% | 98% | 98% | 100% | 99% | 98% | 98% | 100% | 97% | 95% | 99% | 98% |

Results show number of samples sequenced harboring a given *K13* allele in the designated country and year.

[a]Smith SJ, et al. 2018. Efficacy of artemisinin-based combination therapies and prevalence of molecular markers associated with artemisinin, piperaquine and sulfadoxine-pyrimethamine resistance in Sierra Leone. Acta Trop 185: 363-370. doi: 10.1016/j.actatropica.2018.06.016.

[b]Vachot-Ganée L, et al. 2018. A novel field-based molecular assay to detect validated artemisinin-resistant *k13* mutants. Malar J 17: 175. doi: 10.1186/s12936-018-2329-y.

[c]Nzoumbou-Boko R, et al. 2020. Molecular assessment of kelch13 non-synonymous mutations in *Plasmodium falciparum* isolates from Central African Republic (2017–2019). Malar J 19: 191. https://doi.org/10.1186/s12936-020-03264-y

[d]Li J, et al. 2016. Limited artemisinin resistance-associated polymorphisms in *Plasmodium falciparum* K13-propeller and PfATPase6 gene isolated from Bioko Island, Equatorial Guinea. Int J Parasitol Drugs Drug Resist 6: 54-59. doi: 10.1016/j.ijpddr.2015.11.002.

[e]Kakolwa MA, et al. 2018. Efficacy and safety of artemisinin-based combination therapy, and molecular markers for artemisinin and piperaquine resistance in Mainland Tanzania. Malar J 17: 369. doi: 10.1186/s12936-018-2524-x.

[f]Uwimana A, et al. 2020. Emergence and clonal expansion of *in vitro* artemisinin-resistant *Plasmodium falciparum kelch13* R561H mutant parasites in Rwanda. Nat Med 26:1602-

[g]Warsame M, et al. 2019. High therapeutic efficacy of artemether–lumefantrine and dihydroartemisinin–piperaquine for the treatment of uncomplicated *falciparum* malaria in Somalia. Malar J 18: 231. doi: 10.1186/s12936-019-2864-1.

**Table S2. Geographic origin and drug resistance genotypes of *Plasmodium falciparum* clinical isolates and reference lines employed in this study.**

| Parasite | Sanger ID | Original ID | Edited? (parent)[a] | Provider | Geographic origin or source | Year | *K13* | *pfcrt*[b] | *mdr1* | *mdr1* copy number | *dhfr*[c] | *dhps* | *fd* | *mdr2*[d] | *arps10* | *ap-2μ* | *ubp1* |
|---|---|---|---|---|---|---|---|---|---|---|---|---|---|---|---|---|---|
| 3D7$^{WT}$ | -- | 3D7 clone A10 | no | D. Goldberg | Africa | 1981 | WT | WT | WT | 1 | WT | WT | D193Y | WT | WT | WT | WT |
| F32$^{WT}$ | -- | -- | no | F. Benoit-Vical | Tanzania | 1982 | WT | WT | WT | 1 | WT | S436A | WT | S208N/F423Y/I492V | WT | S160N | WT |
| UG659$^{WT}$ | -- | -- | no | P. Rosenthal | Uganda | 2007 | WT | GB4 | Y184F/Y927N | 1 | triple | K540E | WT | S208N/F423Y/I492V | WT | WT | E1528D |
| UG815$^{WT}$ | -- | -- | no | P. Rosenthal | Uganda | 2008 | WT | GB4 | N86Y/D1246Y | 1 | triple | K540E | WT | S208N | WT | WT | WT |
| Dd2$^{WT}$ | -- | Dd2 clone B2 | no | MR4 | Indochina | 1980 | WT | Dd2 | N86Y | 3 | triple | S436F/A613S | D193Y | quadruple | WT | WT | WT |
| Dd2$^{I543T}$ | -- | -- | yes (Dd2) | This lab | Straimer *et al.* | 2015 | I543T | Dd2 | N86Y | 3 | triple | S436F/A613S | D193Y | quadruple | WT | WT | WT |
| Dd2$^{C580Y}$ | -- | -- | yes (Dd2) | This lab | Straimer *et al.* | 2015 | C580Y | Dd2 | N86Y | 3 | triple | S436F/A613S | D193Y | quadruple | WT | WT | WT |
| Cam3.II | PH0306-C | RF 967 | no | R. Fairhurst | Cambodia | 2010 | R539T | Dd2 | Y184F | 1 | triple | S436A/K540E | D193Y | quadruple | V127M/D128H | WT | WT |
| Cam3.II$^{WT}$ | -- | Cam3.II$^{rev}$ | yes (Cam3.II) | This lab | Straimer *et al.* | 2015 | WT | Dd2 | Y184F | 1 | triple | S436A/K540E | D193Y | quadruple | V127M/D128H | WT | WT |
| Cam3.II$^{C580Y}$ | -- | -- | yes (Cam3.II) | This lab | Straimer *et al.* | 2015 | C580Y | Dd2 | Y184F | 1 | triple | S436A/K540E | D193Y | quadruple | V127M/D128H | WT | WT |
| CamWT | PH0164-C | RF 915 | no | R. Fairhurst | Cambodia | 2010 | WT | Dd2 | WT | 1 | triple | S436A/K540E | WT | triple | V127M/D128H | WT | WT |
| CamWT$^{C580Y}$ | -- | -- | yes (CamWT) | This lab | Straimer *et al.* | 2015 | C580Y | Dd2 | WT | 1 | triple | S436A/K540E | WT | triple | V127M/D128H | WT | WT |
| RF7$^{C580Y}$ | PH1008-C | 163-KH1-001RNE | no | R. Fairhurst | Cambodia | 2012 | C580Y | Dd2 + M343L | Y184F | 1 | quadruple | K540N/A581G | D193Y | quadruple | V127M/D128H | WT | WT |
| Thai1$^{WT}$ | -- | TA32A2A4 | no | T. Anderson | Thailand | 2003 | WT | Dd2 | N86Y | 1 | double | K540E/A581G | D193Y | quadruple | V127M/D128H | WT | WT |
| Thai2$^{WT}$ | -- | TA50A2B2 | no | T. Anderson | Thailand | 2004 | WT | Dd2 | F1226Y | 2 | quadruple | K540E/A581G | D193Y | quadruple | V127M/D128H | WT | R3138H |
| Thai3$^{WT}$ | -- | TA85R1 | no | T. Anderson | Thailand | 2003 | WT | Dd2 | F1226Y | 1 | triple | S436A/K540E | D193Y | triple | WT | WT | WT |
| Thai4$^{WT}$ | -- | TA86A3 | no | T. Anderson | Thailand | 2003 | WT | Dd2 | F1226Y | 1 | double | K540E/A581G | D193Y | quadruple | D128Y | WT | WT |
| Thai5$^{WT}$ | -- | NHP-01334-6B | no | T. Anderson | Thailand | 2011 | WT | Dd2 | F1226Y | 1 | quadruple | K540E/A581G | D193Y | quadruple | V127M/D128H | WT | WT |
| Thai6$^{E252Q}$ | -- | NHP4076 | no | T. Anderson | Thailand | 2008 | E252Q | Dd2 | F1226Y | 2 | quadruple | K540N/A581G | D193Y | quadruple | V127M/D128H | WT | WT |
| Thai7$^{E252Q}$ | -- | NHP4673 | no | T. Anderson | Thailand | 2010 | E252Q | Dd2 | WT | 2 | triple | K540N/A581G | D193Y | quadruple | V127M/D128H | WT | WT |

[a]Gene-edited parasite lines were generated by zinc-finger nuclease editing and were previously reported in Straimer J, et al. 2015. K13-propeller mutations confer artemisinin resistance in plasmodium falciparum clinical isolates. Science 347:428-431. doi: 10.1126/science.1260867.

[b]Dd2 *pfcrt*: M74I/N75E/K76T/A220S/Q271E/N326S/I356T/R371I; GB4 *pfcrt:* M74I/N75E/K76T/A220S/Q271E/R371I.

[c]Double *dhfr* mutant: C95R/S108N; Triple *dhfr* mutant: N51I/C59R/S108N; Quadruple *dhfr* mutant: N51I/C59R/S108N/I164L.

[d]Triple *mdr2* mutant: S208N/G299D/F423Y; quadruple *mdr2* mutant: S208N/G299D/F423Y/T484I.

*K13*, Kelch13, PF3D7_1343700; *pfcrt*, *Plasmodium falciparum* chloroquine resistance transporter, PF3D7_0709000; *mdr1*, multi-drug resistance protein 1, PF3D7_0523000; *dhfr*, dihydrofolate reductase, PF3D7_0417200; *dhps,* dihydropteroate synthase, PF3D7_0810800; *fd*, ferredoxin, PF3D7_1318100; *mdr2*, multi-drug resistance protein 2, PF3D7_1447900; *arps10*, apicoplast ribosomal protein S10, PF3D7_1460900.1; *ap-2μ*, AP-2 complex subunit mu, PF3D7_1218300; *ubp1*, ubiquitin binding protein 1, PF3D7_0104300.

**Table S3. Ring-stage survival (RSA) assay data for edited parasites and controls (African strains).**

| Parasite | Parent | K13 | Editing method | RSA survival[a] | | | |
|---|---|---|---|---|---|---|---|
| | | | | Mean | SEM | N[b] | *P* value[c] |
| 3D7$^{WT}$ | -- | WT | -- | 1.4 | 0.3 | 4 | -- |
| 3D7$^{ctrl}$ | 3D7 | WT + bsm | CRISPR-Cas9 | 1.3 | 0.2 | 3 | 0.78 (ns) |
| 3D7$^{R561H}$ | 3D7 | R561H | CRISPR-Cas9 | 6.6 | 0.3 | 3 | <0.0001 (∗∗∗∗) |
| 3D7$^{M579I}$ | 3D7 | M579I | CRISPR-Cas9 | 4.8 | 0.6 | 3 | 0.0023 (∗∗) |
| 3D7$^{C580Y}$ | 3D7 | C580Y | CRISPR-Cas9 | 4.8 | 0.4 | 4 | 0.0006 (∗∗∗) |
| F32$^{WT}$ | -- | WT | -- | 0.3 | 0.04 | 7 | -- |
| F32$^{R561H}$ | F32 | R561H | CRISPR-Cas9 | 0.4 | 0.16 | 3 | 0.21 (ns) |
| F32$^{M579I}$ | F32 | M579I | CRISPR-Cas9 | 0.5 | 0.09 | 6 | 0.043 (∗) |
| F32$^{C580Y}$ | F32 | C580Y | CRISPR-Cas9 | 0.3 | 0.04 | 8 | 0.93 |
| UG659$^{WT}$ | -- | WT | -- | 1.0 | 0.1 | 6 | -- |
| UG659$^{M579I}$ | UG659 | M579I | CRISPR-Cas9 | 6.3 | 0.7 | 6 | <0.0001 (∗∗∗∗) |
| UG659$^{C580Y}$ | UG659 | C580Y | CRISPR-Cas9 | 4.7 | 1.0 | 6 | 0.0046 (∗∗) |
| UG815$^{WT}$ | -- | WT | -- | 1.3 | 0.2 | 7 | -- |
| UG815$^{M579I}$ clone 1 | UG815 | M579I | CRISPR-Cas9 | 11.9 | 1.0 | 7 | <0.0001 (∗∗∗∗) |
| UG815$^{M579I}$ clone 2 | UG815 | M579I | CRISPR-Cas9 | 11.8 | 0.8 | 7 | <0.0001 (∗∗∗∗) |
| UG815$^{C580Y}$ | UG815 | C580Y | CRISPR-Cas9 | 11.8 | 1.5 | 7 | <0.0001 (∗∗∗∗) |
| Dd2$^{WT}$ | -- | WT | -- | 0.6 | 0.1 | 13 | -- |
| Dd2$^{M579I}$ | Dd2 | M579I | CRISPR-Cas9 | 4.0 | 0.5 | 10 | <0.0001 (∗∗∗∗) |
| Dd2$^{C580Y}$ | Dd2 | C580Y | ZFN | 4.7 | 0.4 | 9 | <0.0001 (∗∗∗∗) |

[a]RSA survival values represent the percentage of parasites surviving a 6 h pulse of 700nM dihydroartemisinin, calculated relative to DMSO mock-treated controls. Parasitemias were determined 66 h following drug treatment.

[b]N, number of independent experiments, each with technical duplicates.

[c]*P* value calculated for relative to the respective wild-type parental line. *P* value determined by unpaired *t* test. ns, not significant. ∗ $P$<0.05; ∗∗ $P$<0.01, ∗∗∗ $P$<0.001, ∗∗∗∗ $P$<0.0001.

bsm, binding-site mutations; SEM, standard error of the mean; WT, wild-type.

**Table S4. Fitness assay data (fraction of K13 mutant parasites of total WT and mutant) for edited African parasite lines.**

| | 3D7 vs. 3D7$^{M579I}$ | | | 3D7 vs. 3D7$^{C580Y}$ | | | F32 vs. 3D7$^{M579I}$ | | | F32 vs. F32$^{C580Y}$ | | | UG659 vs. UG659$^{M579I}$ | | | UG659 vs. UG659$^{C580Y}$ | | | UG815 vs. UG815$^{M579I}$ | | | UG815 vs. UG815$^{C580Y}$ | | |
|---|---|---|---|---|---|---|---|---|---|---|---|---|---|---|---|---|---|---|---|---|---|---|---|---|
| Day | Mean[a] | SEM | N[b] | Mean | SEM | N | Mean | SEM | N | Mean | SEM | N | Mean | SEM | N | Mean | SEM | N | Mean | SEM | N | Mean | SEM | N |
| 0[c] | 0.50 | 0.00 | 5 | 0.50 | 0.00 | 5 | 0.50 | 0.00 | 5 | 0.50 | 0.00 | 5 | 0.50 | 0.00 | 2 | 0.50 | 0.00 | 2 | 0.50 | 0.00 | 2 | 0.50 | 0.00 | 2 |
| 4 | 0.40 | 0.04 | 5 | 0.36 | 0.04 | 5 | 0.49 | 0.01 | 5 | 0.60 | 0.08 | 5 | 0.44 | 0.01 | 2 | 0.51 | 0.07 | 2 | 0.27 | 0.04 | 2 | 0.55 | 0.21 | 2 |
| 8 | 0.54 | 0.12 | 5 | 0.52 | 0.13 | 5 | 0.47 | 0.03 | 5 | 0.49 | 0.10 | 5 | 0.43 | 0.04 | 2 | 0.50 | 0.05 | 2 | 0.19 | 0.00 | 2 | 0.40 | 0.16 | 2 |
| 12 | 0.34 | 0.04 | 5 | 0.33 | 0.05 | 5 | 0.44 | 0.01 | 5 | 0.47 | 0.04 | 5 | 0.41 | 0.01 | 2 | 0.49 | 0.07 | 2 | 0.18 | 0.01 | 2 | 0.29 | 0.17 | 2 |
| 16 | 0.28 | 0.05 | 5 | 0.31 | 0.03 | 5 | 0.42 | 0.01 | 5 | 0.41 | 0.05 | 5 | 0.36 | 0.03 | 2 | 0.47 | 0.05 | 2 | 0.15 | 0.07 | 2 | 0.18 | 0.14 | 2 |
| 20 | 0.29 | 0.05 | 5 | 0.36 | 0.06 | 5 | 0.34 | 0.04 | 5 | 0.35 | 0.06 | 5 | 0.34 | 0.00 | 2 | 0.43 | 0.02 | 2 | 0.11 | 0.06 | 2 | 0.24 | n/a | 1 |
| 24 | 0.19 | 0.06 | 5 | 0.23 | 0.04 | 5 | 0.30 | 0.03 | 5 | 0.36 | 0.06 | 5 | 0.31 | 0.02 | 2 | 0.41 | 0.04 | 2 | 0.10 | 0.06 | 2 | 0.21 | n/a | 1 |
| 28 | 0.17 | 0.09 | 5 | 0.28 | 0.06 | 5 | 0.29 | 0.03 | 5 | 0.36 | 0.06 | 5 | 0.26 | 0.00 | 2 | 0.40 | 0.04 | 2 | 0.09 | 0.01 | 2 | 0.18 | n/a | 1 |
| 32 | 0.17 | 0.07 | 5 | 0.17 | 0.02 | 5 | 0.26 | 0.04 | 5 | 0.34 | 0.07 | 5 | 0.19 | 0.01 | 2 | 0.38 | 0.08 | 2 | 0.07 | 0.04 | 2 | 0.18 | n/a | 1 |
| 36 | 0.17 | 0.07 | 4 | 0.11 | 0.04 | 4 | 0.28 | 0.04 | 4 | 0.36 | 0.08 | 4 | 0.2192 | n/a | 1 | 0.42 | 0.00 | 1 | 0.05 | 0.02 | 2 | 0.09 | 0.11 | 2 |
| 40 | 0.15 | 0.09 | 4 | 0.10 | 0.01 | 4 | 0.19 | 0.07 | 4 | 0.25 | 0.10 | 3 | 0.1867 | n/a | 1 | 0.43 | 0.00 | 1 | 0.05 | 0.05 | 2 | 0.09 | 0.08 | 2 |

[a]Fraction of K13 mutant parasites determined by allelic descrimination via TaqMan qPCR.

[b]N, number of independent experiments, each performed in duplicate.

[c]Percent mutant parasites normalized to 50% on Day 0 for all assays.

n/a, not applicable. SEM, standard error of the mean.

**Table S5. Distribution of *K13* alleles over time in Cambodia (2001-2017).**

| Years of sample collection | 2001-02 | | | | 2004-05 | | | | 2006-07 | | | | 2008-09 | | | | 2010-11 | | | | 2012-13 | | | | 2014-15 | | | | 2016-17 | | | | |
|---|---|---|---|---|---|---|---|---|---|---|---|---|---|---|---|---|---|---|---|---|---|---|---|---|---|---|---|---|---|---|---|---|---|
| K13 | W | N | E | S | W | N | E | S | W | N | E | S | W | N | E | S | W | N | E | S | W | N | E | S | W | N | E | S | W | N | E | S | Total |
| WT | 58 | 25 | 66 | n/a | 30 | 27 | 30 | n/a | 9 | 48 | 22 | n/a | 19 | n/a | n/a | n/a | 46 | 110 | 104 | 4 | 12 | 59 | 225 | 13 | 18 | 31 | 74 | 2 | 0 | 68 | 93 | 1 | 1194 |
| G449A | 0 | 0 | 0 | n/a | 0 | 0 | 0 | n/a | 0 | 0 | 0 | n/a | 2 | n/a | n/a | n/a | 0 | 0 | 0 | 0 | 0 | 0 | 0 | 0 | 0 | 0 | 0 | 0 | 0 | 0 | 0 | 0 | 2 |
| N458Y | 0 | 0 | 0 | n/a | 0 | 0 | 0 | n/a | 0 | 0 | 0 | n/a | 0 | n/a | n/a | n/a | 2 | 0 | 0 | 1 | 0 | 0 | 0 | 0 | 0 | 0 | 0 | 0 | 0 | 0 | 0 | 0 | 3 |
| C469F | 0 | 0 | 0 | n/a | 0 | 0 | 0 | n/a | 0 | 0 | 0 | n/a | 0 | n/a | n/a | n/a | 0 | 0 | 0 | 0 | 0 | 0 | 0 | 0 | 0 | 0 | 1 | 0 | 0 | 0 | 0 | 0 | 1 |
| T474I | 0 | 0 | 1 | n/a | 0 | 0 | 0 | n/a | 0 | 1 | 0 | n/a | 0 | n/a | n/a | n/a | 0 | 0 | 0 | 0 | 0 | 0 | 0 | 0 | 0 | 0 | 0 | 0 | 0 | 0 | 0 | 0 | 2 |
| A481V | 0 | 0 | 0 | n/a | 1 | 0 | 0 | n/a | 1 | 0 | 0 | n/a | 2 | n/a | n/a | n/a | 0 | 0 | 0 | 0 | 0 | 0 | 0 | 0 | 0 | 0 | 0 | 0 | 0 | 0 | 0 | 0 | 4 |
| N489D | 0 | 0 | 0 | n/a | 0 | 0 | 0 | n/a | 0 | 0 | 0 | n/a | 0 | n/a | n/a | n/a | 0 | 0 | 0 | 0 | 0 | 0 | 0 | 0 | 0 | 2 | 0 | 0 | 0 | 0 | 0 | 0 | 2 |
| Y493H | 6 | 0 | 0 | n/a | 0 | 0 | 0 | n/a | 9 | 0 | 0 | n/a | 17 | n/a | n/a | n/a | 18 | 21 | 1 | 0 | 0 | 5 | 0 | 0 | 5 | 1 | 0 | 0 | 0 | 0 | 0 | 0 | 83 |
| I543T | 0 | 0 | 0 | n/a | 0 | 0 | 0 | n/a | 0 | 0 | 0 | n/a | 0 | n/a | n/a | n/a | 1 | 0 | 3 | 0 | 0 | 0 | 0 | 0 | 0 | 0 | 0 | 0 | 0 | 0 | 0 | 0 | 4 |
| G533S | 0 | 0 | 1 | n/a | 0 | 0 | 0 | n/a | 0 | 0 | 0 | n/a | 0 | n/a | n/a | n/a | 0 | 0 | 0 | 0 | 0 | 0 | 0 | 0 | 0 | 0 | 0 | 0 | 0 | 0 | 0 | 0 | 1 |
| N537D | 0 | 0 | 0 | n/a | 0 | 0 | 0 | n/a | 0 | 0 | 0 | n/a | 0 | n/a | n/a | n/a | 0 | 0 | 0 | 0 | 0 | 0 | 0 | 0 | 0 | 1 | 0 | 0 | 0 | 0 | 0 | 0 | 1 |
| R539T | 8 | 1 | 0 | n/a | 5 | 0 | 0 | n/a | 7 | 0 | 0 | n/a | 18 | n/a | n/a | n/a | 15 | 21 | 0 | 2 | 1 | 6 | 1 | 0 | 1 | 0 | 0 | 0 | 0 | 0 | 1 | 0 | 87 |
| P553L | 0 | 0 | 0 | n/a | 0 | 0 | 0 | n/a | 0 | 0 | 0 | n/a | 0 | n/a | n/a | n/a | 0 | 0 | 0 | 1 | 0 | 0 | 1 | 0 | 0 | 0 | 2 | 0 | 0 | 0 | 1 | 0 | 5 |
| R561H | 8 | 0 | 0 | n/a | 1 | 0 | 0 | n/a | 0 | 0 | 0 | n/a | 0 | n/a | n/a | n/a | 0 | 0 | 0 | 0 | 0 | 0 | 0 | 0 | 0 | 0 | 0 | 0 | 0 | 0 | 0 | 0 | 9 |
| V568G | 0 | 0 | 0 | n/a | 0 | 0 | 0 | n/a | 0 | 0 | 0 | n/a | 0 | n/a | n/a | n/a | 0 | 0 | 2 | 0 | 0 | 0 | 0 | 0 | 0 | 0 | 0 | 0 | 0 | 0 | 0 | 0 | 2 |
| P574L | 0 | 0 | 1 | n/a | 3 | 0 | 0 | n/a | 0 | 0 | 0 | n/a | 0 | n/a | n/a | n/a | 0 | 0 | 0 | 0 | 0 | 0 | 0 | 0 | 0 | 0 | 0 | 0 | 0 | 0 | 0 | 0 | 4 |
| C580Y | 23 | 0 | 0 | n/a | 41 | 0 | 0 | n/a | 39 | 0 | 0 | n/a | 47 | n/a | n/a | n/a | 191 | 90 | 50 | 11 | 180 | 42 | 16 | 37 | 189 | 156 | 71 | 7 | 33 | 207 | 405 | 80 | 1915 |
| D584V | 0 | 0 | 0 | n/a | 0 | 0 | 0 | n/a | 0 | 0 | 0 | n/a | 4 | n/a | n/a | n/a | 0 | 0 | 0 | 0 | 0 | 0 | 1 | 0 | 0 | 0 | 0 | 0 | 0 | 0 | 0 | 0 | 5 |
| K610R | 0 | 0 | 0 | n/a | 0 | 0 | 0 | n/a | 0 | 0 | 0 | n/a | 0 | n/a | n/a | n/a | 0 | 0 | 0 | 0 | 0 | 0 | 0 | 0 | 0 | 1 | 0 | 0 | 0 | 1 | 0 | 0 | 2 |
| A626E | 0 | 0 | 0 | n/a | 0 | 0 | 0 | n/a | 0 | 0 | 0 | n/a | 0 | n/a | n/a | n/a | 0 | 0 | 0 | 0 | 0 | 0 | 0 | 0 | 0 | 0 | 1 | 0 | 0 | 0 | 0 | 0 | 1 |
| Total | 103 | 26 | 69 | n/a | 81 | 27 | 30 | n/a | 65 | 49 | 22 | n/a | 109 | n/a | n/a | n/a | 273 | 242 | 160 | 19 | 193 | 112 | 244 | 50 | 213 | 192 | 149 | 9 | 33 | 276 | 500 | 81 | 3327 |
| % WT | 56% | 96% | 96% | n/a | 37% | 100% | 100% | n/a | 14% | 98% | 100% | n/a | 17% | n/a | n/a | n/a | 17% | 45% | 65% | 21% | 6% | 53% | 92% | 26% | 8% | 16% | 50% | 22% | 0% | 25% | 19% | 1% | 36% |
| % C580Y | 22% | 0% | 0% | n/a | 51% | 0% | 0% | n/a | 60% | 0% | 0% | n/a | 43% | n/a | n/a | n/a | 70% | 37% | 31% | 58% | 93% | 38% | 7% | 74% | 89% | 81% | 48% | 78% | 100% | 75% | 81% | 99% | 58% |

Results show number of samples sequenced harboring a given *K13* allele in each of four regions of Cambodia during a given span of time. W, Western; N, Northern; E, Eastern; S, Southern.

n/a, not available; WT, wild-type.

**Table S6. Ring-stage survival (RSA) assay data for edited parasites and controls (Southeast Asian strains).**

| Parasite | Parent | *K13* | Editing method | RSA survival[a] | | | |
|---|---|---|---|---|---|---|---|
| | | | | Mean | SEM | N[b] | *P* value[c] |
| Dd2[WT] | -- | WT | -- | 0.6 | 0.1 | 13 | -- |
| Dd2[ctrl] | Dd2 | WT + bsm[d] | CRISPR-Cas9 | 0.9 | 0.1 | 5 | 0.16 (ns) |
| Dd2[E252Q] | Dd2 | E252Q | ZFN | 0.5 | 0.2 | 5 | 0.47 (ns) |
| Dd2[F446I] | Dd2 | F446I | CRISPR-Cas9 | 2.0 | 0.4 | 6 | 0.0004 (***) |
| Dd2[P553L] | Dd2 | P553L | CRISPR-Cas9 | 4.6 | 0.7 | 11 | <0.0001 (****) |
| Dd2[R561H] | Dd2 | R561H | CRISPR-Cas9 | 4.3 | 0.4 | 7 | <0.0001 (****) |
| Dd2[P574L] | Dd2 | P574L | CRISPR-Cas9 | 2.1 | 0.3 | 8 | <0.0001 (****) |
| Dd2[C580Y] | Dd2 | C580Y | ZFN | 4.7 | 0.4 | 9 | <0.0001 (****) |
| Dd2[R539T] | Dd2 | R539T | ZFN | 20.0 | | 1 | <0.0001 (****) |
| Cam3.II[WT] | Cam3.II | WT + bsm[e] | ZFN | 1.5 | 0.5 | 5 | -- |
| Cam3.II[ctrl] | Cam3.II[WT] | WT + bsm[d] | CRISPR-Cas9 | 1.9 | 0.1 | 2 | 0.72 (ns) |
| Cam3.II[E252Q] | Cam3.II[WT] | E252Q | CRISPR-Cas9 | 1.0 | 0.1 | | 0.58 (ns) |
| Cam3.II[F446I] | Cam3.II[WT] | F446I | CRISPR-Cas9 | 1.9 | 0.8 | 4 | 0.73 (ns) |
| Cam3.II[P553L] | Cam3.II[WT] | P553L | CRISPR-Cas9 | 2.8 | 0.4 | 3 | 0.14 (ns) |
| Cam3.II[P574L] | Cam3.II[WT] | P574L | CRISPR-Cas9 | 2.7 | 1.2 | 5 | 0.38 (ns) |
| Cam3.II[R561H] | Cam3.II[WT] | R561H | CRISPR-Cas9 | 13.2 | 1.3 | 6 | <0.0001 (****) |
| Cam3.II[C580Y] | Cam3.II | C580Y | ZFN | 10.0 | | 1 | <0.0001 (****) |
| Cam3.II[R539T] | -- | R539T | -- | 20.4 | 1.3 | 3 | <0.0001 (****) |
| Thai1 | -- | WT | -- | 1.0 | 0.2 | 7 | -- |
| Thai1[E252Q] | Thai1 | E252Q | ZFN | 0.4 | 0.1 | 3 | 0.09 (ns) |
| Thai1[R561H] | Thai1 | R561H | CRISPR-Cas9 | 7.2 | 1.5 | 4 | 0.001 (**) |
| Thai1[C580Y] | Thai1 | C580Y | ZFN | 2.1 | 0.3 | 4 | 0.02 (ns) |
| Thai2 | -- | WT | -- | 1.5 | 0.6 | 3 | -- |
| Thai2[R561H] | Thai2 | R561H | ZFN | 14.5 | 1.7 | 5 | 0.0014 (**) |
| Thai2[C580Y] | Thai2 | C580Y | ZFN | 11.5 | 1.6 | 4 | 0.004 (**) |
| Thai3 | -- | WT | -- | 0.6 | 0.06 | 3 | -- |
| Thai3[E252Q] | Thai3 | E252Q | ZFN | 1.1 | 0.4 | 3 | 0.33 (ns) |
| Thai4 | -- | WT | -- | 0.4 | 0.1 | 3 | -- |
| Thai4[C580Y] | Thai4 | C580Y | ZFN | 15.4 | 1.0 | 3 | 0.0001 (***) |
| Thai5 | -- | WT | -- | 1.2 | 0.2 | 3 | -- |
| Thai5[E252Q] | Thai5 | E252Q | ZFN | 1.4 | 0.1 | 3 | 0.33 (ns) |
| Thai5[R561H] | Thai5 | R561H | ZFN | 12.7 | 1.4 | 4 | 0.0013 (**) |
| Thai5[C580Y] | Thai5 | C580Y | ZFN | 7.6 | 1.2 | 4 | 0.0071 (**) |
| Thai6[E252Q] | -- | E252Q | -- | 2.7 | 0.3 | 4 | -- |
| Thai7[E252Q] | -- | E252Q | -- | 5.1 | 0.8 | 4 | -- |

[a]RSA survival values represent the percentage of parasites surviving a 6 h pulse of 700nM dihydroartemisinin, calculated relative to DMSO mock-treated controls. Parasitemias were determined 66 h following drug treatment.

[b]N, number of independent experiments, each with technical duplicates.

[c]*P* value calculated for relative to the respective wild-type parental line. *P* value determined by unpaired *t* test. ns, not significant. * $P<0.05$; ** $P<0.01$, *** $P<0.001$, **** $P<0.0001$.

[d]The Dd2[ctrl] and Cam3.II[ctrl] lines harbor silent binding-site mutations at the CRISPR-Cas9 cleavage site.

[e]The Cam3.II[WT] line harbors silent binding-site mutations at the zinc-finger nuclease cleavage site.

bsm, binding-site mutations; SEM, standard error of the mean; WT, wild-type; ZFN, zinc-finger nuclease.

**Table S7. Fitness assay data (percent eGFP+ parasites) for edited Dd2 parasites and parental control.**

| | Dd2$^{WT}$ | | | Dd2$^{bsm}$ | | | Dd2$^{E252Q}$ | | | Dd2$^{F446I}$ | | | Dd2$^{P553L}$ | | | Dd2$^{R561H}$ | | | Dd2$^{P574L}$ | | | Dd2$^{C580Y}$ | | |
|---|---|---|---|---|---|---|---|---|---|---|---|---|---|---|---|---|---|---|---|---|---|---|---|---|
| Day | Mean[a] | SEM | N[b] | Mean | SEM | N | Mean | SEM | N | Mean | SEM | N | Mean | SEM | N | Mean | SEM | N | Mean | SEM | N | Mean | SEM | N |
| 0 | 48.9 | 1.4 | 3 | 49.9 | 1.3 | 3 | 48.2 | 4.2 | 3 | 50.2 | 1.2 | 3 | 50.8 | 3.6 | 3 | 47.1 | 3.0 | 3 | 45.2 | 0.7 | 3 | 50.3 | 1.4 | 3 |
| 2 | 40.0 | 3.7 | 3 | 41.9 | 2.8 | 3 | 43.8 | 1.9 | 3 | 42.6 | 7.6 | 3 | 41.6 | 4.5 | 3 | 43.0 | 4.5 | 3 | 39.5 | 5.8 | 3 | 46.2 | 4.3 | 3 |
| 4 | 34.6 | 0.7 | 2 | 38.6 | 1.3 | 2 | 44.5 | 4.5 | 2 | 37.0 | 4.4 | 2 | 39.0 | 2.4 | 2 | 39.4 | 1.5 | 2 | 34.9 | 5.3 | 2 | 42.8 | 0.3 | 2 |
| 6 | 32.1 | 0.3 | 2 | 38.2 | 1.4 | 2 | 45.0 | 4.2 | 2 | 36.3 | 3.7 | 2 | 37.6 | 0.3 | 2 | 39.6 | 2.3 | 2 | 33.6 | 3.0 | 2 | 39.2 | 3.2 | 2 |
| 8 | 25.5 | 1.3 | 2 | 33.1 | 0.3 | 2 | 40.5 | 6.8 | 2 | 32.5 | 0.3 | 2 | 27.1 | 0.7 | 2 | 35.9 | 4.4 | 2 | 29.4 | 1.8 | 2 | 30.7 | 3.6 | 2 |
| 10 | 22.4 | -- | 1 | 32.0 | -- | 1 | 28.8 | -- | 1 | 28.5 | -- | 1 | 23.5 | -- | 1 | 33.7 | -- | 1 | 24.8 | -- | 1 | 24.5 | -- | 1 |
| 12 | 15.3 | 1.1 | 2 | 24.8 | 2.1 | 2 | 25.8 | 0.3 | 2 | 22.0 | 1.5 | 2 | 20.1 | 0.2 | 2 | 27.3 | 1.1 | 2 | 19.1 | 0.5 | 2 | 18.0 | 0.8 | 2 |
| 14 | 14.3 | 1.9 | 3 | 21.6 | 0.5 | 3 | 28.3 | 4.6 | 3 | 18.8 | 0.7 | 3 | 16.3 | 1.7 | 3 | 25.1 | 0.4 | 3 | 15.7 | 1.6 | 3 | 17.1 | 2.0 | 3 |
| 16 | 14.1 | 3.2 | 3 | 24.6 | 2.1 | 3 | 29.5 | 5.5 | 3 | 19.6 | 2.5 | 3 | 17.3 | 3.5 | 3 | 26.5 | 4.1 | 3 | 17.1 | 2.4 | 3 | 15.4 | 4.5 | 3 |
| 18 | 8.9 | 1.1 | 2 | 16.8 | 1.8 | 2 | 23.0 | 5.8 | 2 | 14.0 | 1.2 | 2 | 11.5 | 2.1 | 2 | 21.4 | 3.9 | 2 | 10.6 | 1.3 | 2 | 11.7 | 2.7 | 2 |
| 20 | 7.0 | 1.0 | 3 | 13.9 | 1.8 | 3 | 17.7 | 3.5 | 3 | 11.0 | 1.9 | 3 | 9.0 | 1.6 | 3 | 15.7 | 2.9 | 3 | 8.5 | 2.5 | 3 | 6.6 | 1.4 | 3 |

[a]Percent eGFP+ parasites determined by flow cytometry as a fraction of total (MitoTracker+) parasite population.

[b]N, number of independent experiments, each performed in triplicate.

bsm; binding-site mutations; SEM, standard error of the mean; WT, wild-type.

**Table S8. Ring-stage survival (RSA) assay data for *fd* and *mdr2* edited parasites and parental controls.**

| | | | | | | RSA survival[b] | | | |
|---|---|---|---|---|---|---|---|---|---|
| Parasite | Parent | *K13* | *fd* | *mdr2*[a] | Editing method (locus) | Mean | SEM | N[c] | *P* value[d] |
| CamWT$^{C580Y}$ | CamWT | C580Y | WT | T484 | ZFN *(K13)* | 11.0 | 1.7 | 2 | -- |
| CamWT$^{C580Y\ fd\ D193Y}$ | CamWT$^{C580Y}$ | C580Y | D193Y | T484 | CRISPR-Cas9 *(fd)* | 12.5 | 1.7 | 2 | 0.5798 (ns) |
| RF7$^{C580Y}$ | -- | C580Y | D193Y | T484I | -- | 27.1 | 1.5 | 3 | -- |
| RF7$^{fd\ ctrl}$ | RF7$^{C580Y}$ | C580Y | D193Y | T484I | CRISPR-Cas9 *(fd)* | 27.2 | 2.3 | 3 | 0.9823 (ns) |
| RF7$^{fd\ rev}$ | RF7$^{C580Y}$ | C580Y | WT (rev) | T484I | CRISPR-Cas9 *(fd)* | 26.7 | 2.7 | 3 | 0.9151 (ns) |
| Cam3.II$^{C580Y}$ | Cam3.II | C580Y | D193Y | T484I | ZFN *(K13)* | 18.5 | 1.7 | 3 | -- |
| Cam3.II$^{C580Y\ mdr2\ ctrl}$ | Cam3.II$^{C580Y}$ | C580Y | D193Y | T484I + bsm | CRISPR-Cas9 *(mdr2)* | 21.2 | 1.9 | 3 | 0.3457 (ns) |
| Cam3.II$^{C580Y\ mdr2\ rev}$ | Cam3.II$^{C580Y}$ | C580Y | D193Y | T484 (rev) | CRISPR-Cas9 *(mdr2)* | 21.8 | 3.3 | 3 | 0.4235 (ns) |

[a]All parasites tested carry the S208N/G299D/F423Y mutations in *mdr2* and as such are not designated as wild-type (WT).

[b]RSA survival values represent the percentage of parasites surviving a 6 h pulse of 700nM dihydroartemisinin, calculated relative to DMSO mock-treated controls. Parasitemias were determined 66 h following drug treatment.

[c]N, number of independent experiments, each with technical duplicates.

[d]*P* value calculated relative to the respective parental line. Student's *t* test. ns, not significant.

bsm, binding-site mutations; rev, revertant (denotes a line that has been reverted to wild-type at the given position by gene editing); SEM, standard error of the mean; WT, wild-type.

**Table S9. Oligonucleotides used in this study.**

| Name | Nucleotide sequence (5'-3') | Description | Lab name |
|---|---|---|---|
| p1 | TATTACACATAGCTGATGATCTAG | *K13* gRNA fwd | p6558 |
| p2 | AAACCTAGATCATCAGCTATGTGT | *K13* gRNA rev | p6287 |
| p3 | GTGACGTCGATTGATATTAATGTTGGTGGAGC | *K13* CRISPR/Cas9 donor amplification fwd | p3984 |
| p4 | CCGCATATGGTGCAAACGGAGTGACCAAATCTGGG | *K13* CRISPR/Cas9 donor amplification rev | p3986 |
| p5 | GAATACGCCAAGATCATCAGCTATGTGTGTTGCTTTTGATAATAAAATTTATGTCATTGG | SDM *K13* bsm fwd | p6090 |
| p6 | GCAACACACATAGCTGATGATCTTGGCGTATTCAAAGGTGCCACCTCTACCC | SDM *K13* bsm rev | p6091 |
| p7 | GATAAGAAAGAACTACAAATGGAAAGAGTACGATTGTACAAAGAATTAGAAAACCG | SDM *K13* E252Q fwd | p5174 |
| p8 | CGTACTCTTTCCATTTGTAGTTCTTTCTTATCTTTAAATAATTTATCTTTTTCTCG | SDM *K13* E252Q rev | p5175 |
| p9 | CCCATTAGTAATTTGTATAGGTGGATTTGATGGTGTAGAATATTTAAATTCGATGG | SDM *K13* F446I fwd | p4895 |
| p10 | CCACCTATACAAATTACTAATGGGAATGGTAAAAATTTAATACCATAAAATTCTGC | SDM *K13* F446I fwd | p4896 |
| p11 | GATGGCTCTTCTATTATACTGAATGTAGAAGCATATGATCATCGTATGAAAGCATGG | SDM *K13* P553L fwd | p6086 |
| p12 | TCATATGCTTCTACATTCAGTATAATAGAAGAGCCATCATATCCCCC | SDM *K13* P553L rev | p6087 |
| p13 | GCATATGATCATCATATGAAAGCATGGGTAGAGGTGGCACCTTTGAATACCCC | SDM *K13* R561H fwd | p5176 |
| p14 | CCCATGCTTTCATATGATGATCATATGCTTCTACATTCGGTATAATAGAAGAGCC | SDM *K13* R561H rev | p5177 |
| p15 | GAATACGCTAAGATCATCAGCTATGTGTGTTGCTTTTGATAATAAAATTTATGTCATTGG | SDM *K13* P574L + bsm fwd | p6092 |
| p16 | GCAACACACATAGCTGATGATCTTAGCGTATTCAAAGGTGCCACCTCTACCC | SDM *K13* P574L + bsm rev | p6093 |
| p17 | GAATACGCCAAGATCATCAGCTATATGTGTTGCTTTTGATAATAAAATTTATGTCATTGG | SDM *K13* M579I + bsm fwd | p6013 |
| p18 | GCAACACATATAGCTGATGATCTTGGCGTATTCAAAGGTGCCACCTCTACCC | SDM *K13* M579I + bsm rev | p6014 |
| p19 | GAATACGCCAAGATCATCAGCTATGTATGTTGCTTTTGATAATAAAATTTATGTCATTGG | SDM *K13* C580Y + bsm fwd | p6015 |
| p20 | GCAACATACATAGCTGATGATCTTGGCGTATTCAAAGGTGCCACCTCTACCC | SDM *K13* C580Y + bsm rev | p6016 |
| p21 | GAGGTACCGAGCTCGAATTCGAAACGGAATTAAGTGATGCTAG | *K13* CRISPR/Cas9 donor EcoRI In-Fusion fwd | p6655 |
| p22 | CGAAAAGTGCCACCTGACGTCAAACGGAGTGACCAAATCTGGG | *K13* CRISPR/Cas9 donor AatII In-Fusion rev | p6656 |
| p23 | AACATATGTTAAATATTTATTTCTC | CRISPR/Cas9 donor sequencing fwd | p282 |
| p24 | AGGGTTATTGTCTCATGAGCGG | CRISPR/Cas9 donor sequencing fwd | p283 |
| p25 | AAGCACCGACTCGGTGCCAC | gRNA sequencing rev | p35 |
| p26 | GGGAATCTGGTGGTAACAGC | *K13* integration primer fwd (5' end) | p6176 |
| p27 | CGGAGTGACCAAATCTGGGA | *K13* integration primer rev (3' end) | p6175 |
| p28 | GGTATTAAATTTTTACCATTCCCATTAGTATTTTGTATAGG | *K13* sequencing fwd (internal) | p4186 |
| p29 | TAAGTATATAATATTGTGTACATGTTATCCTAAATGTTTTAGAGCTAGAA | *fd* gRNA In-Fusion fwd | p6156 |
| p30 | TTCTAGCTCTAAAACATTTAGGATAACATGTACACAATATTATATACTTA | *fd* gRNA In-Fusion rev | p6157 |
| p31 | ACCGAGCTCGAATTCATGAATATTGTAATACTATTGTTAATAC | *fd* CRISPR/Cas9 donor amplification / EcoRI In-Fusion fwd | p6239 |
| p32 | GTGCCACCTGACGTCCTGCCTATAAAACTTTCTCGTC | *fd* CRISPR/Cas9 donor amplification / AatII In-Fusion rev | p6238 |
| p33 | AAATATATTCTTTTGTGTACATGTTATCCTAAGAGTGATTGTGTGATTG | SDM fd bsm fwd | p6429 |
| p34 | CAATCACACAATCACTCTTAGGATAACATGTACACAAAAGAATATATTT | SDM fd bsm rev | p6430 |
| p35 | CAAGGAAGACGAACTACACTACATGTAATTTGTCCTCAATC | SDM *fd* D193Y fwd | p6437 |
| p36 | GATTGAGGACAAATTACATGTAGTGTAGTTCGTCTTCCTTG | SDM *fd* D193Y rev | p6438 |
| p37 | CAATATGAACATAAAGTACAACATTAATATATAGC | CRISPR/Cas9 donor sequencing fwd | p6293 |
| p38 | GGGCGACACGGAAATGTTGAATACTC | CRISPR/Cas9 donor sequencing fwd | p6295 |
| p39 | CCAAAAAGGCTATTGCATCAATAAACAGTTTGTATAAT | *fd* integration primer fwd (5' end) | p6630 |
| p40 | GAAAATGTAATTATATACATGTGCAACCCAAAC | *fd* integration primer rev (3' end) | p6631 |
| p41 | GCTCGGGCCCATGAATATTGTAATACTATTGTTAATAC | *fd* sequencing fwd (internal) | p5714 |
| p42 | TAAGTATATAATATTGTTATTCCGGCAACAATAGAGTTTTAGAGCTAGAA | *mdr2* gRNA In-Fusion fwd | p6720 |
| p43 | TTCTAGCTCTAAAACTCTATTGTTGCCGGAATAACAATATTATATACTTA | *mdr2* gRNA In-Fusion rev | p6721 |
| p44 | ACCGAGCTCGAATTCGTTATGTAATAAAATCAGAAAATCCTC | *mdr2* CRISPR/Cas9 donor amplification / EcoRI In-Fusion fwd | p6716 |
| p45 | GTGCCACCTGACGTCCAAATCACTAATATCAGTAAATGATTTAATAATAG | *mdr2* CRISPR/Cas9 donor amplification / AatII In-Fusion rev | p6717 |
| p46 | TTCTGTTTTAATGTATATTATTCCGGCAACAATCGAAGGATTAATAACATGTATTATAT | SDM *mdr2* bsm fwd | p6732 |
| p47 | ATATAATACATGTTATTAATCCTTCGATTGTTGCCGGAATAATATACATTAAAACAGAA | SDM *mdr2* bsm rev | p6733 |
| p48 | CTGTTTTAATGTATATTATTCCGGCAATTATCGAAGGATTAATAACATG | SDM *mdr2* T484I + bsm fwd | p6869 |
| p49 | CATGTTATTAATCCTTCGATAATTGCCGGAATAATATACATTAAAACAG | SDM *mdr2* T484I + bsm rev | p6870 |
| p50 | GTTAACAGAATTGAATACATTAGGAAAGGTGTATGTATTATCCAGACA | *mdr2* integration primer fwd (5' end) | p6918 |
| p51 | GTACATTATTAAATTCAATACTAACACCGAATTTTTTTTCTTGTGATG | *mdr2* integration primer rev (3' end) | p6919 |
| p52 | CTTCTAATTTTGCTTATTTAAATTTTTTCC | *mdr2* sequencing fwd (internal) | p5778 |

bsm, binding-site mutations; fwd, forward; rev, reverse; SDM, site-directed mutagenesis.

**Table S10. Description of gene-editing plasmids generated in this study.**

| Name | Locus | Mutations | Editing method | Lab name |
|---|---|---|---|---|
| pDC2-cam-coSpCas9-U6-gRNA-K13_bsm-hdhfr | *K13* | *K13* bsm | CRISPR/Cas9 | gs3297 |
| pDC2-cam-coSpCas9-U6-gRNA-K13_F446I-hdhfr | *K13* | *K13* bsm + F446I | CRISPR/Cas9 | gs3298 |
| pDC2-cam-coSpCas9-U6-gRNA-K13_P553L-hdhfr | *K13* | *K13* bsm + P553L | CRISPR/Cas9 | gs3272 |
| pDC2-cam-coSpCas9-U6-gRNA-K13_R561H-hdhfr | *K13* | *K13* bsm + R561H | CRISPR/Cas9 | gs3512 |
| pDC2-cam-coSpCas9-U6-gRNA-K13_P574L-hdhfr | *K13* | *K13* bsm + P574L | CRISPR/Cas9 | gs3273 |
| pDC2-cam-coSpCas9-U6-gRNA-K13_M579I-hdhfr | *K13* | *K13* bsm + M579I[a] | CRISPR/Cas9 | gs3264 |
| pDC2-cam-coSpCas9-U6-gRNA-K13_C580Y-hdhfr | *K13* | *K13* bsm + C580Y | CRISPR/Cas9 | gs3262 |
| pZFN18/20-K13_E252Q-hdhfr | *K13* | *K13* bsm + E252Q | ZFN | gs3509 |
| pZFN18/20-K13_R561H-hdhfr | *K13* | *K13* bsm + R561H | ZFN | gs3510 |
| pDC2-cam-Cas9-U6-gRNA-fd_rev_D193-hdhfr | *fd* | *fd* bsm | CRISPR/Cas9 | gs3123 |
| pDC2-cam-Cas9-U6-gRNA-fd_D193Y-hdhfr | *fd* | *fd* bsm + D193Y | CRISPR/Cas9 | gs3114 |
| pDC2-cam-Cas9-U6-gRNA-mdr2_T484-hdhfr | *mdr2* | *mdr2* bsm[b] | CRISPR/Cas9 | gs3277 |
| pDC2-cam-Cas9-U6-gRNA-PfMdr2_T484I-hdhfr | *mdr2* | *mdr2* bsm + T484I[b] | CRISPR/Cas9 | gs3286 |

[a]*K13* donor sequence harbors both CRISPR/Cas9 and ZFN bsm in addition to the M579I mutation.

[b]*mdr2* donor sequence also harbors the S208N/G299D/F423Y mutations found in all parental lines.

bsm, silent binding-site mutations; ZFN, zinc-finger nuclease.

**Table S11. Real-Time PCR (qPCR) primers and probes.**

| Name | Nucleotide sequence (5'-3') | Description |
|---|---|---|
| p7251 | TCGTATGAAAGCATGGGTAGAG | *K13* qPCR fwd |
| p7252 | CCATTAGTTCCACCAATGACATAAA | *K13* qPCR rev |
| p7253 | FAM-5'-CATCAGCTATGTGTGTTGCT-3'-MGB-EclipseDLP | *K13* WT probe with 6-FAM and MGB-Eclipse |
| p7254 | HEX-5'-ATCATCAGCTATG<u>TAT</u>GTTGCT-3'-MGB-EclipseDLP | *K13* 580Y (mutant) probe with 6-HEX and MGB-Eclipse |
| p7373 | HEX-5'-CATCAGCT<u>ATA</u>TGTGTTGCT-3'-MGB-EclipseDLP | K13 579I (mutant) probe with 6-HEX and MGB-Eclipse |

fwd, forward; rev, reverse; WT, wild-type.
